# Fibroblast growth factor-inducible 14 regulates satellite cell self-renewal and expansion during skeletal muscle repair

**DOI:** 10.1101/2024.10.06.616900

**Authors:** Meiricris Tomaz da Silva, Aniket S. Joshi, Ashok Kumar

## Abstract

Skeletal muscle regeneration in adults is predominantly driven by satellite cells. Loss of satellite cell pool and function leads to skeletal muscle wasting in many conditions and disease states.

Here, we demonstrate that the levels of fibroblast growth factor-inducible 14 (Fn14) are increased in satellite cells after muscle injury. Conditional ablation of Fn14 in Pax7-expressing satellite cells drastically reduces their expansion and skeletal muscle regeneration following injury. Fn14 is required for satellite cell self-renewal and proliferation as well as to prevent precocious differentiation. Targeted deletion of Fn14 inhibits Notch signaling but leads to the spurious activation of STAT3 signaling in regenerating skeletal muscle and in cultured muscle progenitor cells. Silencing of STAT3 improves proliferation and inhibits premature differentiation of Fn14-deficient satellite cells. Furthermore, conditional ablation of Fn14 in satellite cells exacerbates myopathy in the mdx mouse model of Duchenne muscular dystrophy (DMD) whereas its overexpression improves the engraftment of exogenous muscle progenitor cells into the dystrophic muscle of mdx mice. Altogether, our study highlights the crucial role of Fn14 in the regulation of satellite cell fate and function and suggests that Fn14 can be a potential molecular target to improve muscle regeneration in muscular disorders.

## INTRODUCTION

Adult skeletal muscle possesses a robust capacity to regenerate which is predominantly attributed to the presence of muscle stem cells, known as satellite cells (1–3). Under normal conditions, satellite cells reside between the basal lamina and the sarcolemma of myofibers in a reversible quiescent state. Satellite cells express paired box 7 (Pax7) protein, which is essential for their survival, self-renewal, and the orchestration of proper muscle regeneration (4). In response to muscle injury, satellite cells rapidly induce MyoD expression, undergo several rounds of cell division, and eventually differentiate into myoblasts, which fuse with each other or with injured myofibers to accomplish muscle repair (3). While most activated satellite cells differentiate into myoblasts, a small proportion of them repress MyoD expression and returns to quiescence to respond to the next round of muscle injury and regeneration (1, 5, 6). Both extrinsic alteration in muscle niche and intrinsic alteration in cell autonomous regulatory mechanisms in satellite cells, such as reduced asymmetric cell division, myogenic commitment, and senescence are responsible for the loss of skeletal muscle mass in many conditions, including aging and degenerative muscle disorders, such as muscular dystrophy (7–9).

The abundance and myogenic function of satellite cells are regulated through the coordinated activation of multiple signaling pathways. Notch signaling, which is activated by physical interaction between a cell that expresses Notch ligands (Dll1, 4 and Jag1, 2) and a cell that expresses one of the four Notch receptors (Notch1–4), plays a particularly crucial role in the maintenance of the satellite cell pool and the regulation of cell fate decisions (10–13).

Constitutive activation of Notch signaling promotes the self-renewal of satellite cells by upregulating Pax7, whereas its disruption in satellite cells results in impaired muscle regeneration (10, 14–16). In addition to Notch, a few other signaling pathways, such as MAPKs, canonical nuclear factor-kappa B (NF-κB), and the Wnt planar cell polarity (PCP) pathway also regulate the expansion of satellite cells during adult muscle regeneration (17–21). By contrast, other signaling pathways, such as canonical Wnt/β-catenin promote satellite cell myogenic lineage progression by limiting Notch signaling and thus promoting differentiation (12). Recent studies have also demonstrated that deregulation of JAK-STAT signaling is an important contributor to satellite cell dysfunction in aging muscle and dystrophic muscle (22–24) and an interplay between Notch and JAK-STAT signaling is essential for the temporal regulation of satellite cell fate, ensuring proper coordination between activation and differentiation during skeletal muscle regeneration (25).

Satellite cells are activated by various cytokines and growth factors that are released by both myogenic and non-myogenic cells in injured muscle microenvironment (26). TWEAK and its receptor Fn14 constitute a ligand-receptor dyad that activates multiple signaling pathways to regulate inflammation and cellular proliferation and differentiation (27, 28). Intriguingly, a few studies have demonstrated that overexpression of Fn14 is sufficient to activate downstream signaling pathways even in the absence of TWEAK cytokine (27, 29, 30). A prior study showed that muscle regeneration is significantly reduced in global Fn14-knockout (KO) mice potentially due to a diminished inflammatory immune response (31). More recently, we reported that Fn14 promotes myoblast fusion during skeletal muscle regeneration (32). However, the role and mechanisms of action of the Fn14 receptor-mediated signaling in satellite cell homeostasis and function during skeletal muscle regeneration remain unknown. Furthermore, the role of Fn14 in the regulation of satellite cell function in models of degenerative muscle disorders, such as Duchenne muscular dystrophy (DMD) remains completely unknown.

In the present study, using genetic mouse models, we have investigated the role of Fn14 in the regulation of satellite cell function during muscle regeneration. Our results demonstrate that the levels of Fn14 are upregulated in satellite cells following muscle injury. Satellite cell-specific deletion of Fn14 inhibits the self-renewal and proliferation of satellite cells and reduces muscle regeneration in response to acute injury. Our results demonstrate that while targeted ablation of Fn14 inhibits Notch signaling, it leads to spurious activation of STAT3 both *in vivo* and *in vitro* which contributes to satellite cell dysfunction. Finally, our results demonstrate that conditional deletion of Fn14 in satellite cells exacerbates dystrophic phenotype in the mdx mice, which mimic DMD pathology. In contrast, overexpression of Fn14 in exogenous wild type myogenic cells improves their engraftment into the dystrophic muscle of mdx mice following transplantation.

## RESULTS

### Dynamic activation of Fn14 signaling in satellite cells during muscle regeneration

We first investigated how the expression of Fn14 is regulated during muscle regeneration in adult mice. Using a published single cell RNA-seq (scRNA-seq) dataset (GSE143435) (33), we analyzed the gene expression of Fn14 (gene name: *Tnfrsf12a*) in satellite cells at different time points after injury. In this study, the tibialis anterior (TA) muscle of adult C57BL/6J was injured by a single intramuscular injection of notexin. The TA muscle was collected at various time points (0 [uninjured], 2, 5, and 7 days) followed by preparation of single cell suspension and scRNA-seq (33). Our analysis showed that Fn14 is expressed in satellite cells of uninjured TA muscle.

Intriguingly, the mRNA levels of Fn14 were increased in satellite cells at day 2 post-injury, and persisted through days 5 and 7 post-injury. By contrast, the mRNA levels of TWEAK (gene name: *Tnfsf12*) in satellite cells were low in uninjured muscle and did not show any increase after muscle injury (**Figure 1A**).

**FIGURE 1.**
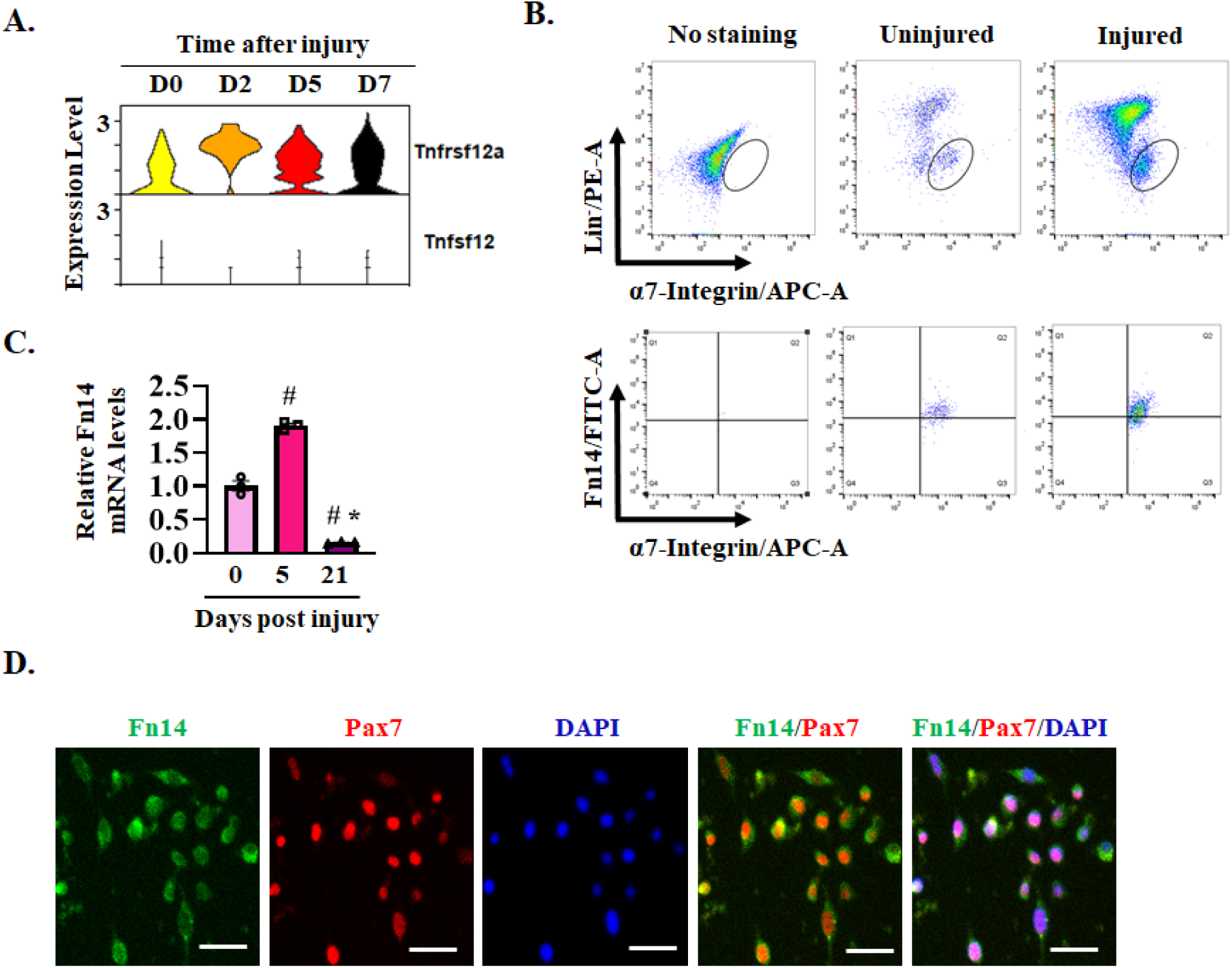
Expression of Fn14 in satellite cells. **(A)** Violin plots showing gene expression of Fn14 (gene name: *Tnfrsf12a*) and TWEAK (gene name: *Tnfsf12*) in satellite cells at different time points after muscle injury in mice analyzed from a publicly available scRNA-seq dataset (GSE143435). **(B)** Single cells isolated from uninjured and injured TA muscle of WT mice were subjected to FACS analysis for the expression of α7-integrin and Fn14 protein. Representative scatter plots of FACS-based analysis demonstrate presence of Fn14^+^ cells amongst α7-integrin^+^ population (lower panel). **(C)** Satellite cells isolated from TA muscle of WT mice at day 0, 5, or 21 post-injury were analyzed for mRNA levels of Fn14 by performing qPCR. n= 3 mice in each group. Data are presented as mean ± SEM and analyzed by one-way ANOVA followed by Tukey’s multiple comparison test. #p ≤ 0.05, values significantly different from uninjured muscle (day 0). *p ≤ 0.05, values significantly different from 5d-injured TA muscle. **(D)** Representative photomicrographs showing expression of Fn14 in Pax7^+^ cells in cultured mouse primary myoblasts. Scale bar: 50 µm.

We also studied the expression of Fn14 in satellite cells of uninjured and regenerating skeletal muscle of mice. To induce regeneration, the right side TA muscle of 8-week-old male C57BL/6J mice was injured by intramuscular injection of 1.2% BaCl_2_ solution whereas the contralateral TA muscle severed as uninjured control. The muscle tissue was collected on day 5 post-injury and Fn14 expression in satellite cells was analyzed by FACS-based approach using a well-established panel of cell surface markers, Lin^-^ (CD31^-^, CD45^-^, Sca-1^-^, TER119^-^) and α7- Integrin^+^ (17). Our analysis showed that Fn14 was present in the α7-Integrin^+^ cells of both uninjured and injured TA muscles of mice **(Figure 1B)**. We also investigated how the expression of Fn14 in satellite cells is regulated on day 21, a time point when muscle regeneration is complete, and satellite cells begin to return to quiescence. For this experiment, the TA muscle of adult C57BL/6J mice was injured for 5 or 21 days, followed by isolation of satellite cells by magnetic-activated cell sorting (MACS) approach and performing qPCR. Results showed that the mRNA levels of Fn14 were significantly higher in satellite cells of 5d-injured TA muscle compared with uninjured muscle. Interestingly, the mRNA levels of Fn14 in satellite cells were significantly lower in 21d-injured TA muscle compared to both uninjured and 5d-injured TA muscle **(Figure 1C)**. These results suggest a transient increase in the expression of Fn14 in satellite cells during muscle regeneration.

Satellite cells are rapidly activated (express MyoD) and transition to the progenitor stage after isolation and culturing. By performing immunostaining for Fn14 and Pax7 (a marker of satellite cells) proteins, we investigated the expression of Fn14 in myoblast cultures established from hindlimb muscle of C57BL/6J mice. Consistent with the scRNA-seq analysis, Fn14 protein was present in Pax7^+^ cells in myoblast cultures **(Figure 1D)**. These results demonstrate that Fn14 is expressed in muscle progenitor cells both in vivo and in vitro.

### Inducible ablation of Fn14 in satellite cells inhibits muscle regeneration in adult mice

We investigated whether the satellite cell-specific deletion of Fn14 affects skeletal muscle regeneration in adult mice. Floxed Fn14 (henceforth Fn14^fl/fl^) mice were crossed with tamoxifen-inducible satellite cell-specific Cre mice (*Pax7-CreERT2*) to generate satellite cell-specific inducible Fn14 knockout mice (i.e., Fn14^fl/fl^;*Pax7-CreERT2,* henceforth Fn14^scKO^) and littermate Fn14^fl/fl^ mice. To induce Fn14 deletion, 7-week old male Fn14^scKO^ mice were given daily intraperitoneal injections of tamoxifen (75 mg/kg body weight) for 4 consecutive days. The mice were kept on a tamoxifen-containing chow for the entire duration of the study. Littermate male Fn14^fl/fl^ were also treated with tamoxifen and served as controls. Next, one side TA muscle of Fn14^fl/fl^ and Fn14^scKO^ mice was injected with 1.2% BaCl_2_ solution, whereas the contralateral uninjured TA muscle served as a control. Muscle regeneration was evaluated on day 5, 14 and 21 post-injury (**Figure 2A**). To confirm the inactivation of Fn14 in satellite cells of Fn14^scKO^ mice, we performed FACS and qPCR analyses. Results showed that the proportion of Fn14^+^/α7-Integrin^+^ cells and the mRNA levels of Fn14 in purified satellite cells were significantly reduced in 5d-injured TA muscle of Fn14^scKO^ mice compared to corresponding muscle of Fn14^fl/fl^ mice (**Supplemental Figure 1, A-C**).

**FIGURE 2.**
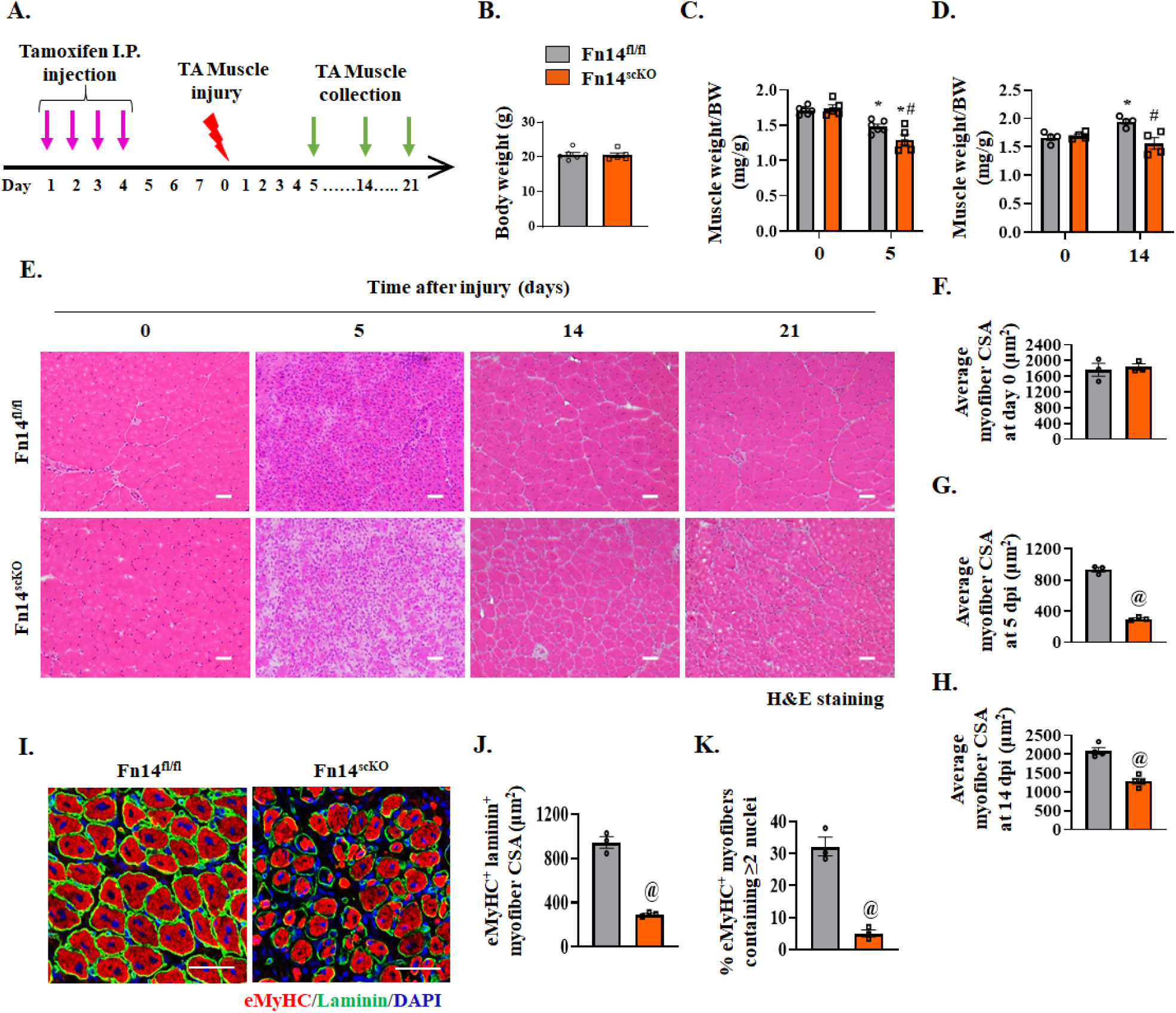
Satellite cell-specific ablation of Fn14 inhibits muscle regeneration. **(A)** Schematic representation of the experimental design indicating tamoxifen regimen for Fn14 deletion, muscle injury, and collection. **(B)** Body weight of Fn14^fl/fl^ and Fn14^scKO^ mice. **(C)** Uninjured and 5d-injured TA muscle wet weight normalized by body weight (BW) of Fn14^fl/fl^ and Fn14^scKO^ mice. **(D)** Uninjured and 14d-injured TA muscle wet weight normalized by BW of Fn14^fl/fl^ and Fn14^scKO^ mice. **(E)** Representative photomicrographs of H&E-stained transverse sections of TA muscle of Fn14^fl/fl^ and Fn14^scKO^ mice at indicated time points after injury. Scale bar: 50 µm. Quantitative analysis of average myofiber cross-sectional area (CSA) in TA muscle of Fn14^fl/fl^ and Fn14^scKO^ mice on **(F)** day 0, **(G)** day 5 and **(H)** day 14 post-injury. **(I)** Representative photomicrographs of transverse sections of 5d-injured TA muscle of Fn14^fl/fl^ and Fn14^scKO^ mice after immunostaining for embryonic isoform of myosin heavy chain (eMyHC) and laminin protein. Nuclei were identified by staining with DAPI. Scale bar: 50 µm. Quantification of **(J)** average CSA of eMyHC^+^ laminin^+^ myofibers, **(K)** percentage of eMyHC^+^ laminin^+^ myofibers containing 2 or more centrally located nuclei in 5d-injured TA muscle of Fn14^fl/fl^ and Fn14^scKO^ mice. n= 3-6 mice in each group. All data are presented as mean ± SEM. *p ≤ 0.05, values significantly different from contralateral uninjured muscle of Fn14^fl/fl^ or Fn14^scKO^ mice; #p ≤ 0.05, values significantly different from corresponding 5d- or 14d-injured TA muscle of Fn14^fl/fl^ mice analyzed by two-way ANOVA followed by Tukey’s multiple comparison test. ^@^p ≤ 0.05, values significantly different from corresponding Fn14^fl/fl^ mice analyzed by unpaired Student *t* test.

There was no significant difference in body weight or wet weight of muscle normalized by body weight between Fn14^fl/fl^ and Fn14^scKO^ mice under naïve conditions (**Figure 2, B-D**).

However, there was a significant decrease in muscle wet weight normalized by body weight in Fn14^scKO^ mice compared to Fn14^fl/fl^ mice on day 5 and 14 post-injury (**Figure 2, C and D**).

Next, transverse sections of TA muscle were generated, followed by performing Hematoxylin and Eosin (H&E) staining and morphometric analysis. There was no significant difference in the myofiber cross-sectional area (CSA) in TA muscle of Fn14^fl/fl^ and Fn14^scKO^ mice in naïve muscle (**Figure 2, E and F**). However, muscle regeneration was significantly impaired in Fn14^scKO^ mice compared to their littermate Fn14^fl/fl^ mice. At both day 5 and 14 post-injury, there was a significant decrease in myofiber CSA of Fn14^scKO^ mice compared to Fn14^fl/fl^ mice (**Figure 2, E, G, and H**). Notably, defects in muscle regeneration (e.g., reduced myofiber CSA and presence of cellular infiltrate) in Fn14^scKO^ mice persisted even on day 21 post-injury (**Figure 2E**). In a separate experiment, we also investigated whether the deletion of only one allele of the Fn14 gene in satellite cells contributes to the observed phenotype in Fn14^scKO^ mice. We compared muscle regeneration defects in male Fn14^fl/wt^;*Pax7-CreERT2* (Fn14^fl/scKO^) mice with male Fn14^scKO^ mice. There was a significant deficit in muscle regeneration in Fn14^scKO^ mice compared to the corresponding Fn14^fl/scKO^ or Fn14^fl/fl^ mice at day 5 post-injury (**Supplemental Figure 1, D-F**).

Deficits in muscle regeneration in Fn14^scKO^ mice were also evidenced by immunostaining of 5d-injured TA muscle sections for embryonic isoform of MyHC (eMyHC) which is expressed in newly formed myofibers (**Figure 2I**). Average CSA of eMyHC^+^ myofibers and the proportion of eMyHC^+^ myofibers containing 2 or more centrally located nuclei were significantly reduced in 5d-injured TA muscle of Fn14^scKO^ mice compared to Fn14^fl/fl^ mice (**Figure 2, J and K**). We also investigated the expression of eMyHC in regenerating myofibers at day 14 and 21 post-injury (**Supplemental Figure 2, A-C)**. There were almost no eMyHC^+^ myofibers in the TA muscle of Fn14^fl/fl^ mice at day 14 or 21 after injury suggesting normal progression of muscle regeneration. In contrast, small-sized eMyHC^+^ myofibers were abundant in the muscle of Fn14^scKO^ mice on day 14 post-injury (**Supplemental Figure 2, A and B)**. The presence of eMyHC^+^ myofibers in TA muscle section of Fn14^scKO^ mice was even prolonged to day 21 post-injury (**Supplemental Figure 2C)**. These results suggest that satellite cell-specific inducible deletion of Fn14 inhibits skeletal muscle regeneration in adult mice.

To understand the role of Fn14 in the regulation of the abundance of satellite cells, uninjured and 5d-injured TA muscle sections of Fn14^fl/fl^ and Fn14^scKO^ mice were immunostained for Pax7 and laminin protein. Nuclei were counterstained with DAPI (**Figure 3A**). There was no significant difference in the number of satellite cells in uninjured TA muscle of Fn14^fl/fl^ and Fn14^scKO^ mice (**Supplemental Figure 3, A and B**). However, the number of satellite cells per unit area was significantly reduced in the TA muscle of Fn14^scKO^ mice compared to Fn14^fl/fl^ mice at day 5 and 14 post-injury (**Figure 3, A-C**). We also performed FACS analysis and quantified the abundance of satellite cells (i.e., Lin^−^ α7-Integrin^+^) in uninjured and 5d-injured TA muscle of Fn14^fl/fl^ and Fn14^scKO^ mice. Consistent with immunohistochemistry results, there was no significant difference in the proportion of satellite cells in uninjured TA muscle of Fn14^fl/fl^ and Fn14^scKO^ mice (**Supplemental Figure 3, C and D**). In contrast, the proportion of satellite cells in Fn14^scKO^ mice was significantly reduced in 5d-injured TA muscle of Fn14^scKO^ mice compared to Fn14^fl/fl^ mice (**Figure 3, D and E**). Moreover, the mRNA levels of Pax7 were also significantly reduced in 5d-injured TA muscle of Fn14^scKO^ mice compared to Fn14^fl/fl^ mice (**Figure 3F**).

**FIGURE 3.**
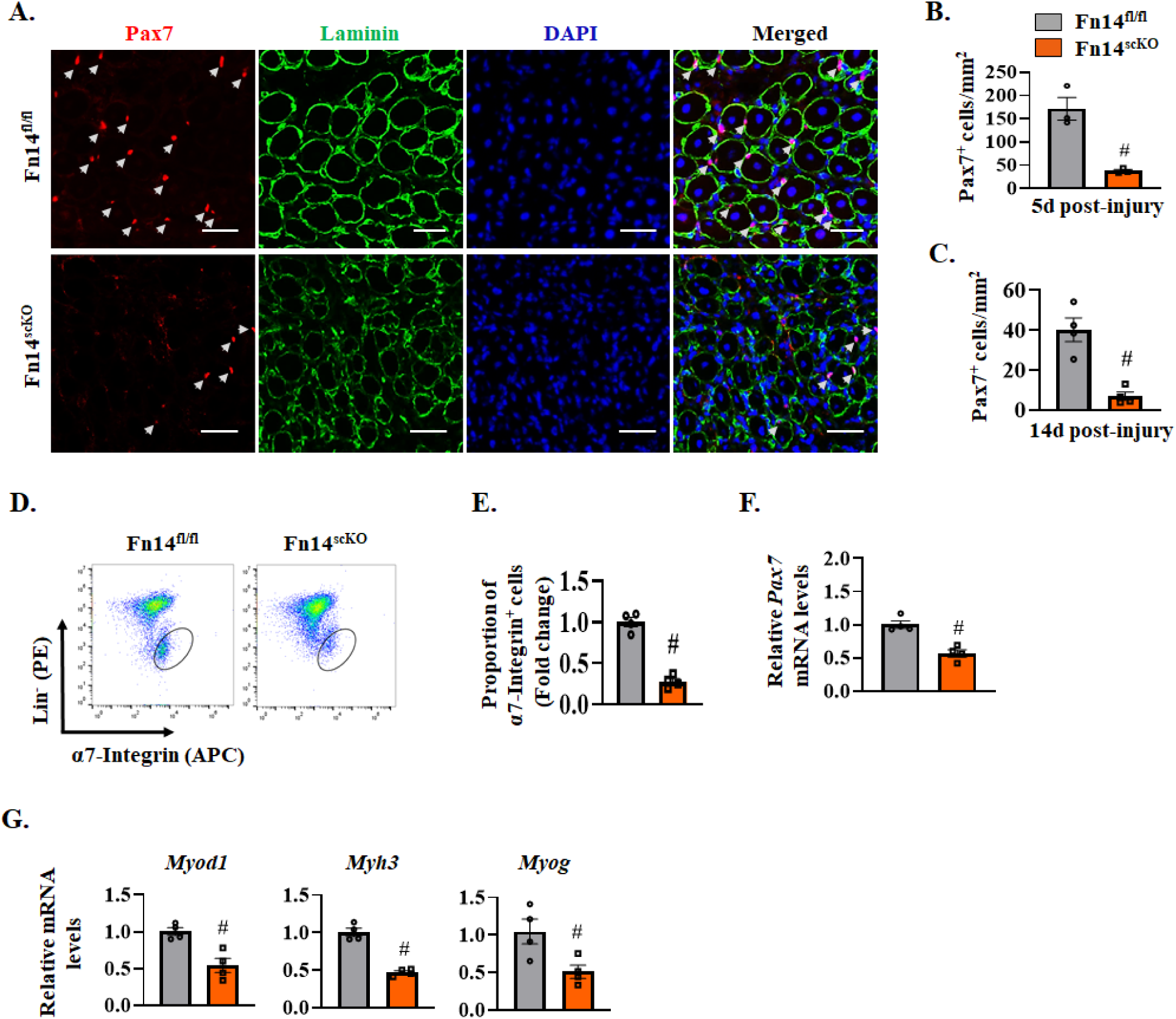
Fn14 regulates satellite cell number in regenerating skeletal muscle. **(A)** Representative photomicrographs of 5d-injured TA muscle sections of Fn14^fl/fl^ and Fn14^scKO^ mice after immunostaining for Pax7 (red) and laminin (green) protein. Nuclei were identified by staining with DAPI. Scale bar: 50 µm. White arrows point to Pax7^+^ satellite cells. **(D)** Average number of Pax7^+^ cells per unit area in **(B)** 5d-injured and **(C)** 14d-injured TA muscle of Fn14^fl/fl^ and Fn14^scKO^ mice. **(D)** Single cell suspensions were isolated from TA muscle of Fn14^fl/fl^ and Fn14^scKO^ mice were subjected to FACS analysis for satellite cells. Representative FACS dot plots demonstrating the percentage of α7-integrin^+^ cells in 5d-injured TA muscle of Fn14^fl/fl^ and Fn14^scKO^ mice. **(E)** Quantification of α7-integrin^+^ satellite cells in 5d-injured TA muscle of Fn14^fl/fl^ and Fn14^scKO^ mice assayed by FACS. Relative mRNA levels of **(E)** *Pax7*, **(F)** *Myod1*, *Myh3* and *Myog* in 5d-injured TA muscle of Fn14^fl/fl^ and Fn14^scKO^ mice assayed by performing qPCR. n= 3-4 mice in each group. All data are presented as mean ± SEM. #p ≤ 0.05, values significantly different from corresponding muscle of Fn14^fl/fl^ mice analyzed by unpaired Student *t* test.

Skeletal muscle regeneration relies on the hierarchical expression of myogenic regulatory factors (MRFs), such as Myf5, MyoD, and myogenin, along with the embryonic isoform of myosin-heavy chain (eMyHC). This sequential expression is crucial for completing the myogenic program and determines the efficiency of muscle repair (1, 34). Our analysis showed that the mRNA levels of *Myod1*, *Myh3* (eMyHC), and *Myog* (myogenin) were significantly reduced in injured TA muscle of Fn14^scKO^ mice compared to Fn14^fl/fl^ mice (**Figure 3G**).

Collectively, these results suggest that targeted inducible deletion of Fn14 reduces the number of satellite cells and attenuates regeneration of injured skeletal muscle in adult mice.

### Fn14 is required for the proliferation of satellite cells

While tamoxifen-inducible approach, using a Cre-ERT2 system, is considered highly efficient for excising floxed alleles, it does not always result in a complete deletion of the target gene due to various factors, including incomplete recombination efficiency that results in a mixed population of cells with varying levels of gene expression. Furthermore, the cells that did not undergo complete gene deletion by tamoxifen-inducible approach may proliferate at a faster rate and overgrow the knockout (KO) cells after a few passages in culture. Therefore, for in vitro experiments, we employed primary myoblast cultures established from skeletal muscle of germline whole body Fn14-KO mice that ensures complete and sustained ablation of Fn14 protein in all myogenic cells. To understand the mechanisms of action of Fn14 in muscle progenitor cells, we performed bulk RNA sequencing (RNA-seq) using primary myoblast cultures established from hindlimb muscle of wild type (WT) and Fn14-KO mice. Functional enrichment analysis of the differentially expressed genes (DEGs) using gene ontology (GO) annotations showed that several gene sets related to cell projection morphogenesis, skeletal system development, muscle cell proliferation, cell-cell signaling and positive regulation of stem cell proliferation were downregulated in Fn14-KO compared with WT cultures. By contrast, the gene sets involved in the regulation of regulation of myoblast and myotube differentiation, negative regulation of cell proliferation, and tyrosine phosphorylation of STAT protein were upregulated in Fn14-KO cultures (**Figure 4A**). Further analysis of DEGs in the RNA-seq experiment showed that gene expression of many molecules associated with cell proliferation and migration were downregulated in Fn14-KO cultures compared to WT cultures (**Figure 4B, Supplemental Figure 4A**). To experimentally validate the role of Fn14 in the proliferation of myogenic cells, WT and Fn14-KO cells were incubated in growth medium for 48 h and pulsed with EdU for 1 h, followed by detection of EdU^+^ cells (**Figure 4C**). There was a significant reduction in the proportion of EdU^+^ cells in Fn14-KO cultures compared to WT cultures (**Figure 4D**). In another experiment, myoblast cultures were also stained with anti-Pax7 and EdU (**Supplemental Figure 4B**) followed by quantification of Pax7^+^/EdU^+^ cells. Results showed that the proportion of Pax7^+^/EdU^+^ cells was significantly reduced in Fn14-KO cultures compared to WT cultures (**Supplemental Figure 4C**). LDH is a stable enzyme present in all cells and released following plasma membrane damage and cell death (35). Our analysis showed that there was no significant difference in the levels of LDH in culture supernatants of WT and Fn14-KO myoblast cultures, suggesting that the deficiency of Fn14 does not affect the survival of myogenic cells (**Supplemental Figure 4D**).

**FIGURE 4.**
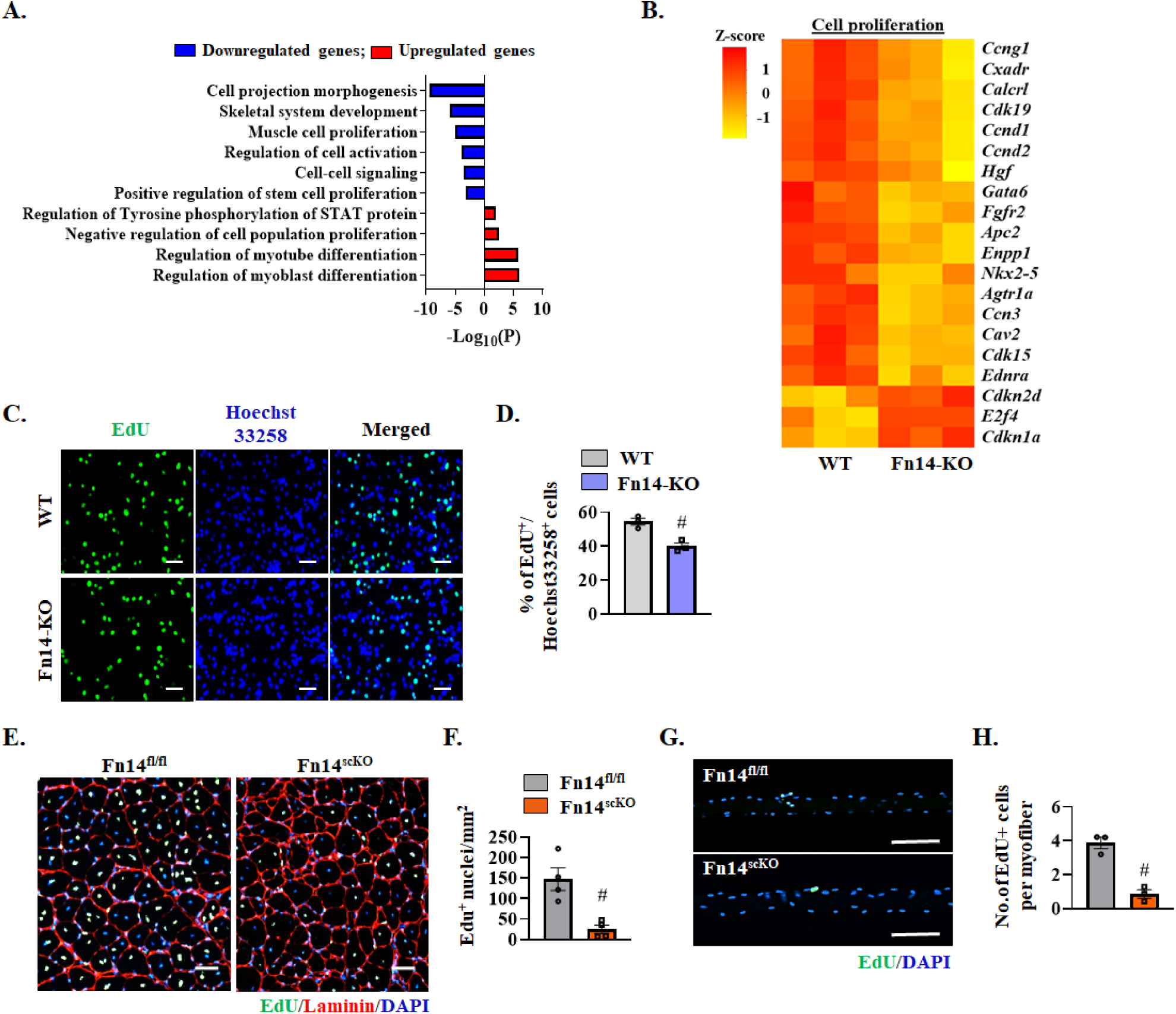
Fn14 mediates activation and proliferation of satellite cells. **(A)** Gene ontology (GO) biological processes associated with downregulated and upregulated genes. **(B)** Heatmap representing selected genes involved in the regulation of cell proliferation in cultured WT and Fn14-KO myogenic cells generated after analysis of RNA-seq dataset. **(C)** WT and Fn14-KO myogenic cultures were pulse-labeled with EdU for 60 min. Representative images of the cultures after detection of EdU and Hoechst staining (nuclei detection). Scale bar: 50 µm. **(D)** Quantification of percentage of EdU^+^ cells in WT and Fn14-KO cultures. n= 3 biological replicates in each group. **(E)** TA muscle of Fn14^fl/fl^ and Fn14^scKO^ mice was injured by intramuscular injection of 1.2% BaCl_2_ solution. After 3 days, the mice were given an intraperitoneal injection of EdU and 11 days later TA muscles were collected, transverse muscle sections were generated and stained to detect EdU, anti-laminin, and nuclei. Representative photomicrographs after EdU, laminin, and DAPI staining are presented here. Scale bar: 50 µm. **(F)** Quantification of the EdU^+^ nuclei per unit area. n= 4 mice in each group. **(G)** Single myofibers were isolated from EDL muscle of Fn14^fl/fl^ and Fn14^scKO^ mice. After 48 h of culturing, the myofibers were pulse-labeled with EdU for 60 min. Representative merged images of EdU^+^ and DAPI-stained myofibers are presented here. Scale bar: 100 μm. **(H)** Quantification of number of EdU^+^ cells per myofiber. n= 3 mice in each group. Analysis was done using 15-20 myofibers for each mouse. n= 3 mice in each group. All data are presented as mean ± SEM. #p ≤ 0.05, values significantly different from WT myoblast, or corresponding muscle of Fn14^fl/fl^ mice, analyzed by unpaired Student *t* test.

We next sought to investigate the role of Fn14 in the proliferation of satellite cells in response to muscle injury in vivo. For this experiment, TA muscle of 8-week old male Fn14^fl/fl^ and Fn14^scKO^ mice was injured by intramuscular injection of 1.2% BaCl_2_ solution. The mice were given a single intraperitoneal injection of EdU at day 3 after injury and the TA muscle was isolated at day 14 post-injury and processed for detection of EdU, DAPI staining, and immunostaining for laminin protein (**Figure 4E**). Quantitative analysis showed that the number of EdU^+^ nuclei per unit area (mm^2^) was significantly reduced in TA muscle of Fn14^scKO^ mice compared to Fn14^fl/fl^ mice (**Figure 4F**). To further validate the role of Fn14 in satellite cell proliferation, we also established single myofiber cultures from extensor digitorum longus (EDL) muscle of male Fn14^fl/fl^ and Fn14^scKO^ mice and the myofiber-associated satellite cells were analyzed after 48 h of culturing. Consistent with in vitro and in vivo results, we found a significant reduction in the number of EdU^+^ cells per myofiber in cultures from Fn14^scKO^ mice compared to Fn14^fl/fl^ mice (**Figure 4, G and H**). Altogether, these results suggest that Fn14 is essential for the proliferation of muscle progenitor cells.

### Fn14 is required for satellite cell self-renewal

In addition to generating sufficient number of myoblasts during skeletal muscle regeneration, activated satellite cells play a crucial role in maintaining their own reserve pool via self-renewal, which is essential for sustaining continuous muscle regeneration throughout the lifespan of the organism (36). To investigate the role of Fn14 in satellite cell self-renewal, we employed a re-injury model, involving two injuries separated by 3-4 weeks (32, 37, 38). After 21 days of the first injury, the TA muscle of 8-week old male Fn14^fl/fl^ and Fn14^scKO^ mice was re-injured using intramuscular injection of BaCl_2_ solution. The TA muscle was isolated on day 5 after the second injury and analyzed by performing H&E staining, Sirius red staining, and immunostaining for Pax7 and laminin protein (**Figure 5, A and D**). There was a significant reduction in the proportion of myofibers containing 2 or more nuclei in TA muscle of Fn14^scKO^ mice compared to Fn14^fl/fl^ mice (**Figure 5B**). Additionally, there was an increase in connective tissue in Fn14^scKO^ mice compared to Fn14^fl/fl^ mice assessed by Sirius red staining (**Figure 5C**). Remarkably, the number of Pax7^+^ cells per unit area (mm^2^) was drastically reduced in TA muscle of Fn14^scKO^ mice compared to Fn14^fl/fl^ mice after double-injury (**Figure 5E**).

**FIGURE 5.**
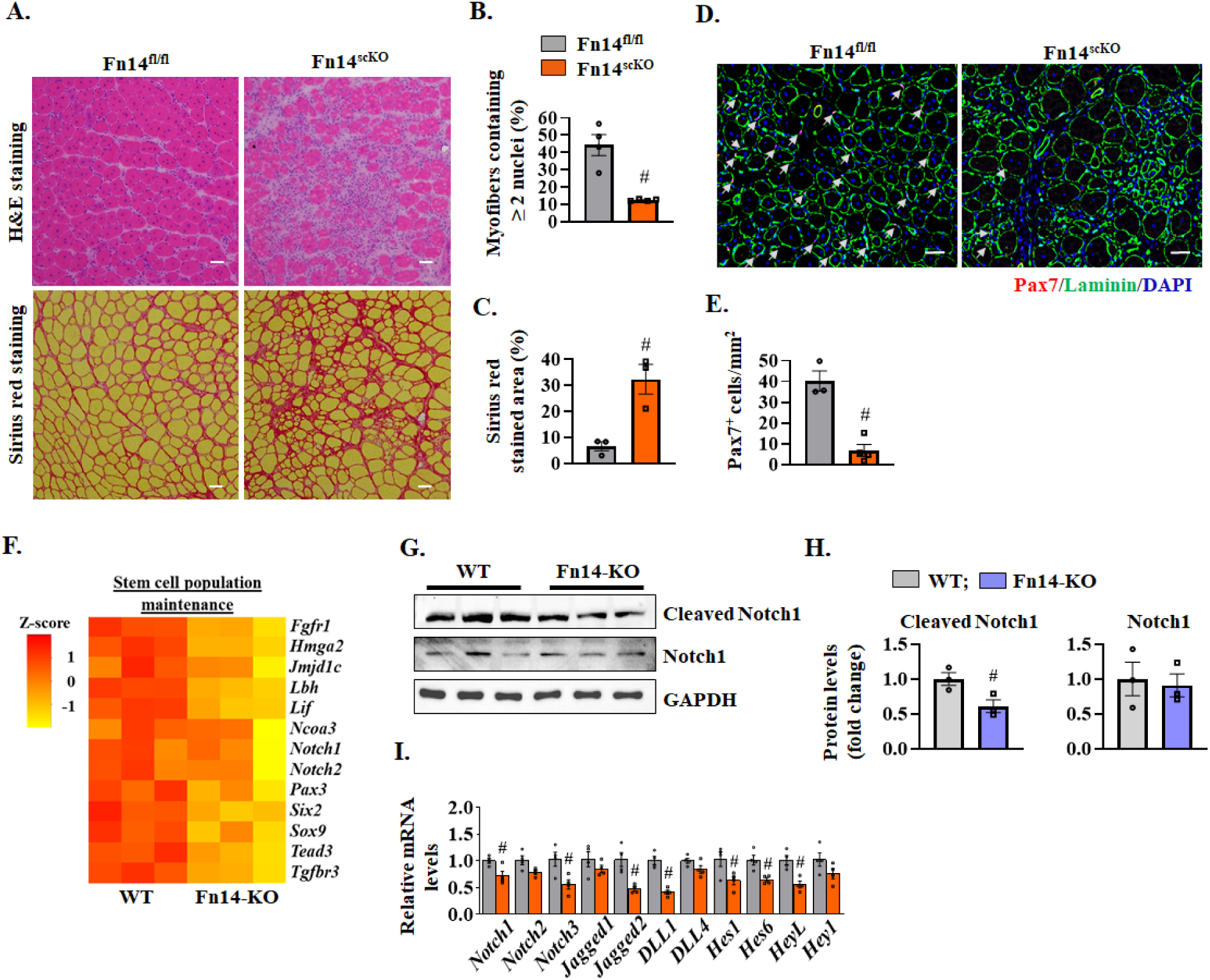
Fn14 promotes satellite cell self-renewal. **(A)** TA muscle of Fn14^fl/fl^ and Fn14^scKO^ mice was injured by intramuscular injection of 1.2% BaCl_2_ solution. After 21 days, the same TA muscle was injured again, and the muscle was isolated 5 days later. Representative photomicrograph of H&E-and Sirius red-stained TA muscle sections. Quantification of **(B)** percentage of myofibers containing two or more centrally located nuclei and **(C)** percentage of total area stained with Sirius red. Scale bar: 50 µm. **(D)** Representative photomicrograph and **(E)** quantification of Pax7^+^ cells per unit area in double injured TA muscle sections of Fn14^fl/fl^ and Fn14^scKO^ mice after immunostaining for Pax7 (red) and laminin (green) protein. Nuclei were identified by staining with DAPI. Scale bar: 50 µm. n= 3-4 mice in each group. **(F)** Heatmap of selected genes associated with stem cell population maintenance in WT and Fn14-KO cultures generated after analysis of RNA-seq dataset. **(G)** Immunoblots and **(H)** densitometry analysis showing levels of cleaved Notch1 and total Notch1 protein in WT and Fn14-KO cultures. n= 3 biological replicates in each group. **(I)** Relative mRNA levels of Notch receptors (*Notch1*, *Notch2*, *Notch3*), Notch ligands (*Jagged1*, *Jagged2*, *Dll1*, and *Dll4*), and Notch targets (*Hes1*, *Hes6*, *Heyl*, *and Hey1*) in 5d-injured TA muscle of Fn14^fl/fl^ and Fn14^scKO^ mice. n= 4 mice in each group. All data are presented as mean ± SEM. #p ≤ 0.05, values significantly different from corresponding muscle of Fn14^fl/fl^ mice analyzed by unpaired Student *t* test.

We next investigated potential mechanisms through which Fn14 improves self-renewal of satellite cells. The GO term analysis of DEGs in RNA-seq experiment revealed a significant downregulation in the gene expression of molecules involved in cell-cell signaling in Fn14-KO cultures (**Figure 4A**). Notch is an important cell-cell signaling pathway that is essential for satellite cell self-renewal and proliferation during regenerative myogenesis (14, 39–41). Indeed, our RNA-seq analysis showed that gene expression of several molecules associated with the maintenance of stem cell population, including components of Notch signaling was downregulated in primary myoblast cultures of Fn14-KO mice compared to WT mice (**Figure 5F**). We also measured the levels of Notch1 protein in WT and Fn14-KO myoblast cultures.

Interestingly, levels of cleaved Notch1 protein were significantly reduced in Fn14-KO cultures compared to WT cultures indicating an inhibition in Notch signaling in Fn14-KO cultures (**Figure 5, G and H**). Furthermore, by performing qPCR assay, we compared the mRNA levels of Notch receptors (*Notch1*, *Notch2*, and *Notch3*), Notch ligands (*Jagged1*, *Jagged2*, *Dll1*, and *Dll4*), and Notch target genes (*Hes1*, *Hes6*, *HeyL*, and *Hey1*) in 5d-injured TA muscle of Fn14^fl/fl^ and Fn14^scKO^ mice. There was a significant reduction in the mRNA levels of *Notch1*, *Notch3*, *Jagged2*, *Dll1*, *Hes1*, *Hes6* and *HeyL* in 5d-injured TA muscle of Fn14^scKO^ mice compared to corresponding muscle of Fn14^fl/fl^ mice (**Figure 5I**). These results suggest that targeted deletion of Fn14 inhibits self-renewal of satellite cells.

### Fn14 prevents precocious differentiation of satellite cells

GO analysis of DEGs in RNA-seq experiment demonstrated significant upregulation of pathways related to myoblast and myotube differentiation (**Figure 4A**). Further analysis of DEGs showed that mRNA levels of several molecules associated with muscle differentiation, including myogenin and Myf6, were significantly upregulated in Fn14-KO cultures compared to WT cultures (**Figure 6A**). Based on the expression of Pax7 and MyoD, the progeny of satellite cell can be classified as self-renewing (Pax7^+^, MyoD^-^), activated/proliferating (Pax7^+^, MyoD^+^), or differentiating (Pax7^-^, MyoD^+^) (38, 42). To directly assess the role of Fn14 in myogenic differentiation, equal number of WT and Fn14-KO primary myoblasts were seeded in culture plates and incubated in growth medium (GM) for 24 h followed by immunostaining for Pax7 and MyoD proteins. Results showed that the proportion of Pax7^+^/MyoD^+^ cells was significantly decreased whereas the proportion of Pax7^-^/MyoD^+^ cells was significantly increased in Fn14-KO cultures compared to WT cultures (**Supplemental Figure 4, E-G**). Furthermore, western blot analysis showed a significant decrease in the protein levels of Pax7 and a drastic increase in the levels of myogenin in Fn14- KO cultures compared to WT cultures (**Figure 6, B and C**).

**FIGURE 6.**
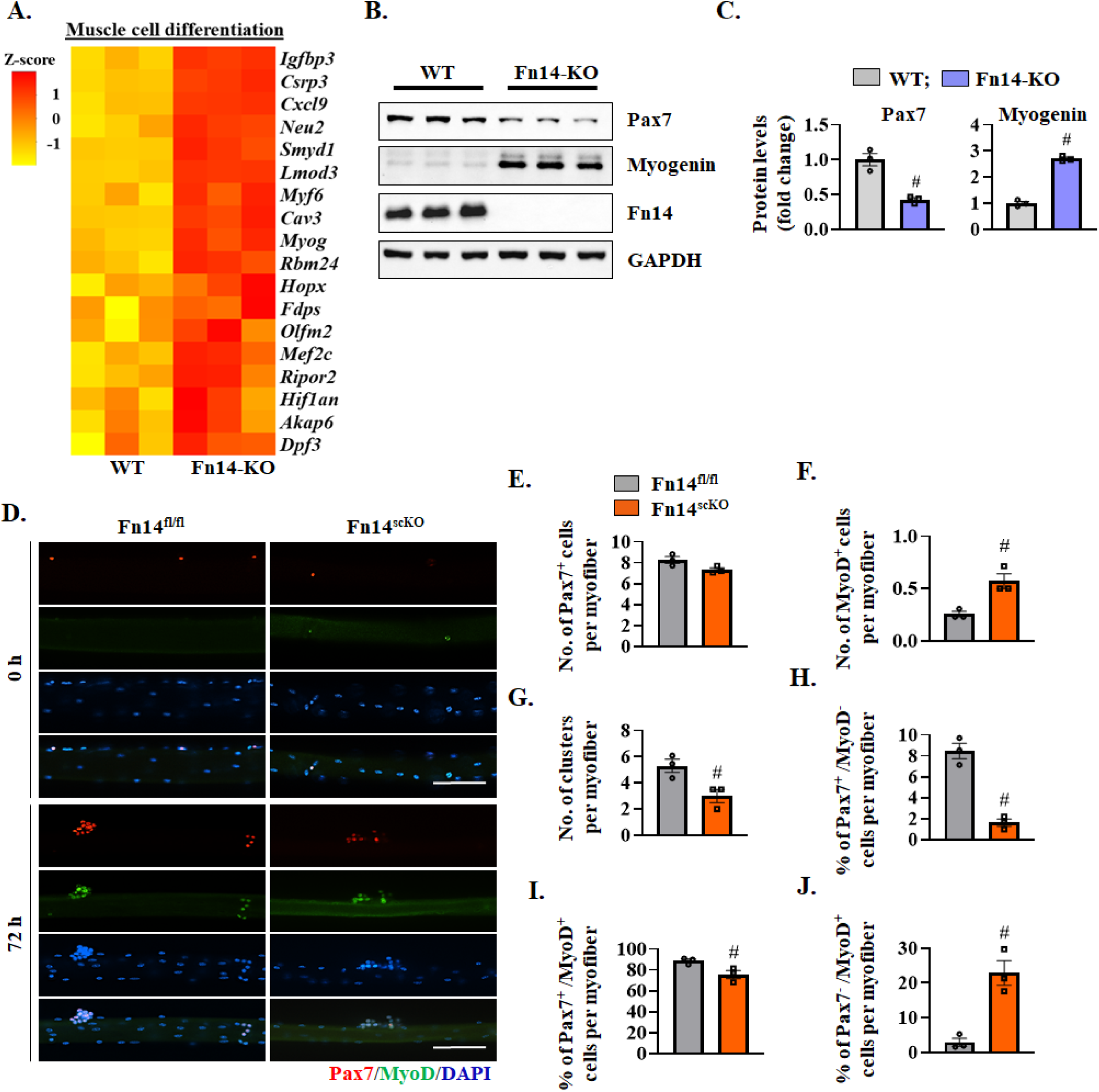
Deletion of Fn14 leads to precocious differentiation of satellite cells. **(A)** Heatmap representing selected genes associated with muscle cell differentiation in WT and Fn14-KO myogenic cultures generated after analysis of RNA-seq dataset. **(B)** Immunoblots and **(C)** densitometry analysis of protein levels of Pax7, myogenin, Fn14, and unrelated protein GAPDH in WT and Fn14-KO cultures. n= 3 biological replicates in each group. **(D)** Single myofibers were isolated from EDL muscle of Fn14^fl/fl^ and Fn14^scKO^ mice and fixed at 0 h or at 72 h of culturing and stained for Pax7 and MyoD protein. Nuclei were counterstained with DAPI. Representative individual-stained and merged images of 0 h and 72 h cultured myofibers from Fn14^fl/fl^ and Fn14^scKO^ mice. Scale bars: 100 μm. Quantification of **(E)** number of Pax7^+^ cells per myofiber, and **(F)** number of MyoD^+^ cells per myofiber at 0 h. Quantification of **(G)** number of clusters per myofiber, percentage of **(H)** Pax7^+^/MyoD^-^ (self-renewing), **(I)** Pax7^+^/MyoD^+^ (activated/proliferating), and **(J)** Pax7^-^/MyoD^+^ (differentiating) cells per myofiber following 72 h of culturing. Analysis was done using 15-20 myofibers for each mouse at each time point. n= 3 mice in each group. All data are presented as mean ± SEM. #p ≤ 0.05, values significantly different from WT myoblast, or corresponding muscle of Fn14^fl/fl^ mice analyzed by unpaired Student *t* test.

We next investigated whether Fn14 plays a role in the self-renewal, proliferation, or differentiation of myofiber-associated satellite cells ex vivo. We established single myofiber cultures from the EDL muscle of male Fn14^fl/fl^ and Fn14^scKO^ mice and performed anti-Pax7 and anti-MyoD at 0 h and 72 h of culturing (**Figure 6D**). While there was no significant difference in the number of Pax7^+^ cells per myofiber, a significant increase in the number of MyoD^+^ cells per myofiber was observed in the freshly isolated myofibers of Fn14^scKO^ mice compared to Fn14^fl/fl^ mice (**Figure 6, E and F**). After 72 h in suspension culture, the number of clusters per myofiber was significantly reduced in myofiber cultures prepared from Fn14^scKO^ mice compared to Fn14^fl/fl^ mice (**Figure 6G**). Importantly, there was a significant decrease in the proportion of Pax7^+^/MyoD^-^ (self-renewing) and Pax7^+^/MyoD^+^ (activated) cells along with a significant increase in the proportion of Pax7^-^/MyoD^+^ (differentiating) cells per myofiber in Fn14^scKO^ cultures compared to Fn14^fl/fl^ cultures (**Figure 6, H-J**). Collectively, these results suggest that the absence of Fn14 leads to premature differentiation of muscle progenitor cells.

### Deletion of Fn14 leads to hyper-activation of STAT signaling

The JAK-STAT signaling plays an important role in the regulation of satellite cell function and muscle regeneration (22, 23). Interestingly, GO term analysis of DEGs from RNA-seq experiment showed a significant upregulation in the molecules involved in the regulation of tyrosine phosphorylation of STAT protein in Fn14-KO cultures (**Figure 4A**). Further analysis of DEGs revealed that gene expression of several positive regulators of the Jak-Stat pathway and target molecules was upregulated whereas gene expression of certain molecules associated with the inhibition of Jak-Stat signaling was downregulated in Fn14-KO cultures compared to WT cultures (**Figure 7A**). To experimentally validate the role of Fn14 in the regulation of Jak-Stat pathway, we first investigated how the satellite cell-specific deletion of Fn14 affects the activation of Jak-Stat signaling in skeletal muscle of mice in response to injury. Western blot analysis showed a dramatic increase in phosphorylation of Stat3 protein at S727 and Y705 residue without having any effect on total levels of Stat3 protein in 5d-injured TA muscle of Fn14^scKO^ mice compared to corresponding muscle of Fn14^fl/fl^ mice (**Figure 7B, Supplemental Figure 5A**). There was also a small but significant increase in the levels of phospo-Stat2 in 5d-injured TA muscle of Fn14^scKO^ mice compared with Fn14^fl/fl^ mice. Interestingly, levels of total Stat2 and Stat5 proteins were also considerably higher in 5d-injured TA muscle of Fn14^scKO^ mice compared with Fn14^fl/fl^ mice. In contrast, there was no difference in the levels of phospho-Stat5 or total Stat3 protein in injured TA muscle of Fn14^fl/fl^ and Fn14^scKO^ mice (**Figure 7B, Supplemental Figure 5A**). Similar to in vivo results, we also found an increase in the levels of phospho-Stat3 protein in Fn14-KO cultures compared to WT cultures of myogenic cells (**Figure 7C, Supplemental Figure 5B**). We next sought to determine whether activated Stat3 has any role in the proliferation and differentiation of Fn14-KO myogenic cells. Primary myoblasts prepared from WT and Fn14-KO mice were transfected with control or Stat3 siRNA. After 24 h of transfection, cellular proliferation was evaluated by pulse labeling of the cells with EdU for 60 min (**Figure 7D**).

**FIGURE 7.**
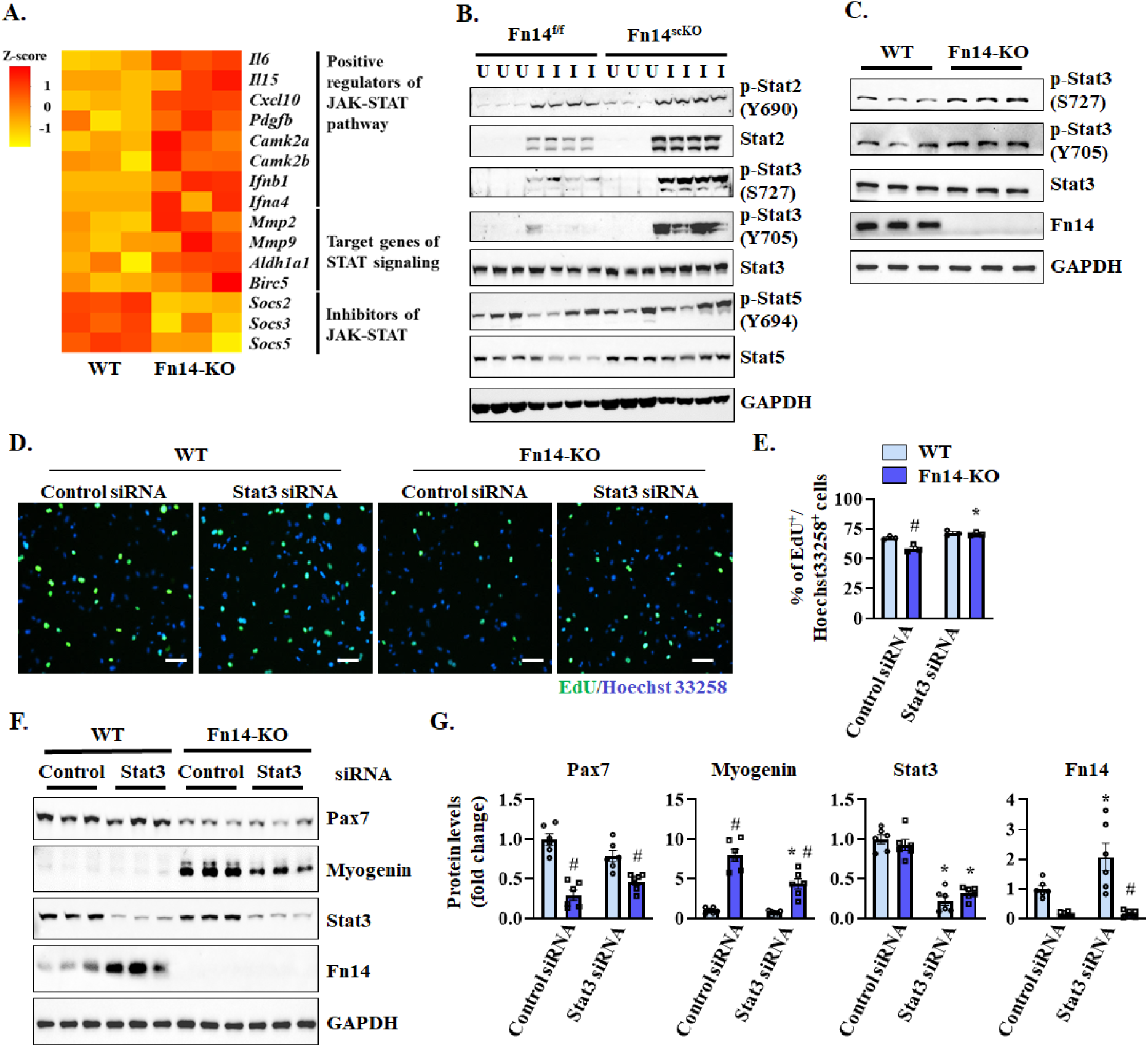
Fn14 regulates STAT signaling in myogenic cells. **(A)** Heatmap of selected genes associated with positive regulation of Jak-Stat pathway, target genes, and inhibitors of Jak-Stat signaling in WT and Fn14-KO cultures generated after analysis of RNA-seq dataset. **(B)** Immunoblots showing protein levels of phosphorylated and total levels of Stat2, Stat3 and Stat5 protein in uninjured and 5d-injured TA muscle of Fn14^fl/fl^ and Fn14^scKO^ mice. n= 3-4 mice in each group. **(C)** Immunoblots showing protein levels of phosphorylated and total Stat3 and Fn14 protein in WT and Fn14-KO cultures. **(D)** WT and Fn14-KO myoblasts were transfected with control and Stat3 siRNA. After 24 h, the cells were pulse-labeled with EdU for 60 min and analyzed for EdU incorporation. Representative photomicrographs are presented here. Scale bar: 100 µm. **(E)** Quantification of percentage of EdU^+^/Hoechst33258^+^ cells in WT and Fn14-KO cultures transfected with control or Stat3 siRNA. n= 3 biological replicates in each group. **(F)** Representative immunoblots and **(G)** densitometry analysis showing protein levels of Pax7, myogenin, Stat3 and Fn14 in WT and Fn14-KO myogenic cultures transfected with control or Stat3 siRNA. n= 6 biological replicates in each group. All data are presented as mean ± SEM. *p ≤ 0.05, values significantly different from corresponding cultures transfected with control siRNA; #p ≤ 0.05, values significantly different from corresponding WT cultures analyzed by two-way ANOVA followed by Tukey’s multiple comparison test.

Results showed that the siRNA-mediated knockdown of Stat3 significantly increased the proportion of EdU^+^ cells in Fn14-KO cultures compared to cultures transfected with control siRNA (**Figure 7E**). By performing Western blot, we also investigated the effect of silencing of Stat3 on the protein levels of Pax7 and myogenin in WT and Fn14-KO cultures. While there was no significant effect on the levels of Pax7 protein, knockdown of Stat3 significantly reduced the levels of myogenin protein in Fn14-KO cultures (**Figure 7, F and G**). Interestingly, knockdown of Stat3 increased levels of Fn14 protein, suggesting a feedback mechanism between Fn14 and Stat3 signaling (**Figure 7, F and G**). These results suggest that the upregulation of Stat3 signaling contributes to reduced proliferation and precocious differentiation of satellite cells.

### Deletion of Fn14 in satellite cells attenuates regeneration and exacerbates myopathy in dystrophic muscle of mdx mice

Dystrophic muscle of mdx and D2.mdx mice (mouse models of DMD) undergo repetitive cycles of degeneration and regeneration, making them a valuable model for studying muscle repair in response to chronic injury (43–45). We first analyzed published scRNA-seq dataset (GSE213925) (46) to determine how the gene expression of Fn14 is regulated in muscle stem cells of dystrophic muscle of mdx and D2.mdx mice. Interestingly, transcript levels of Fn14, TWEAK, and various components of canonical and non-canonical NF-κB signaling were reduced in satellite cells of mdx and D2.mdx mice compared to corresponding WT mice (**Figure 8A**). To understand the role of satellite cell Fn14 signaling in the progression of dystrophic phenotype, we crossed Fn14^scKO^ mice with mdx mice to generate littermate mdx;Fn14^scKO^ and mdx;Fn14^fl/fl^ mice. At the age of 3 weeks, male mdx;Fn14^fl/fl^ and mdx;Fn14^scKO^ mice were given daily i.p. injections of tamoxifen for four consecutive days to induce Cre-mediated recombination and were kept on a tamoxifen-containing chow for the entire duration of the study. The mice were analyzed at the age of 7 weeks. Deletion of Fn14 led to a significant reduction in the size and body weight of mdx;Fn14^scKO^ mice compared to mdx;Fn14^fl/fl^ mice (**Figure 8, B and C**). Moreover, wet weight of TA, gastrocnemius (GA), and quadriceps (QUAD) muscles normalized by body weight was significantly reduced in mdx;Fn14^scKO^ mice compared to mdx;Fn14^fl/fl^ mice (**Figure 8D**). Analysis of H&E-stained GA muscle sections showed signs of exacerbated myopathy marked by a significant decrease in the number of centronucleated fibers (CNF) in mdx; Fn14^scKO^ mice compared to mdx; Fn14^fl/fl^ mice (**Figure 8, E and F**). Moreover, the average CSA of eMyHC^+^ myofibers and the number of Pax7^+^ cells per unit area were also significantly reduced in mdx;Fn14^scKO^ mice compared to mdx;Fn14^fl/fl^ mice (**Figure 8, E, G and H**).

**FIGURE 8.**
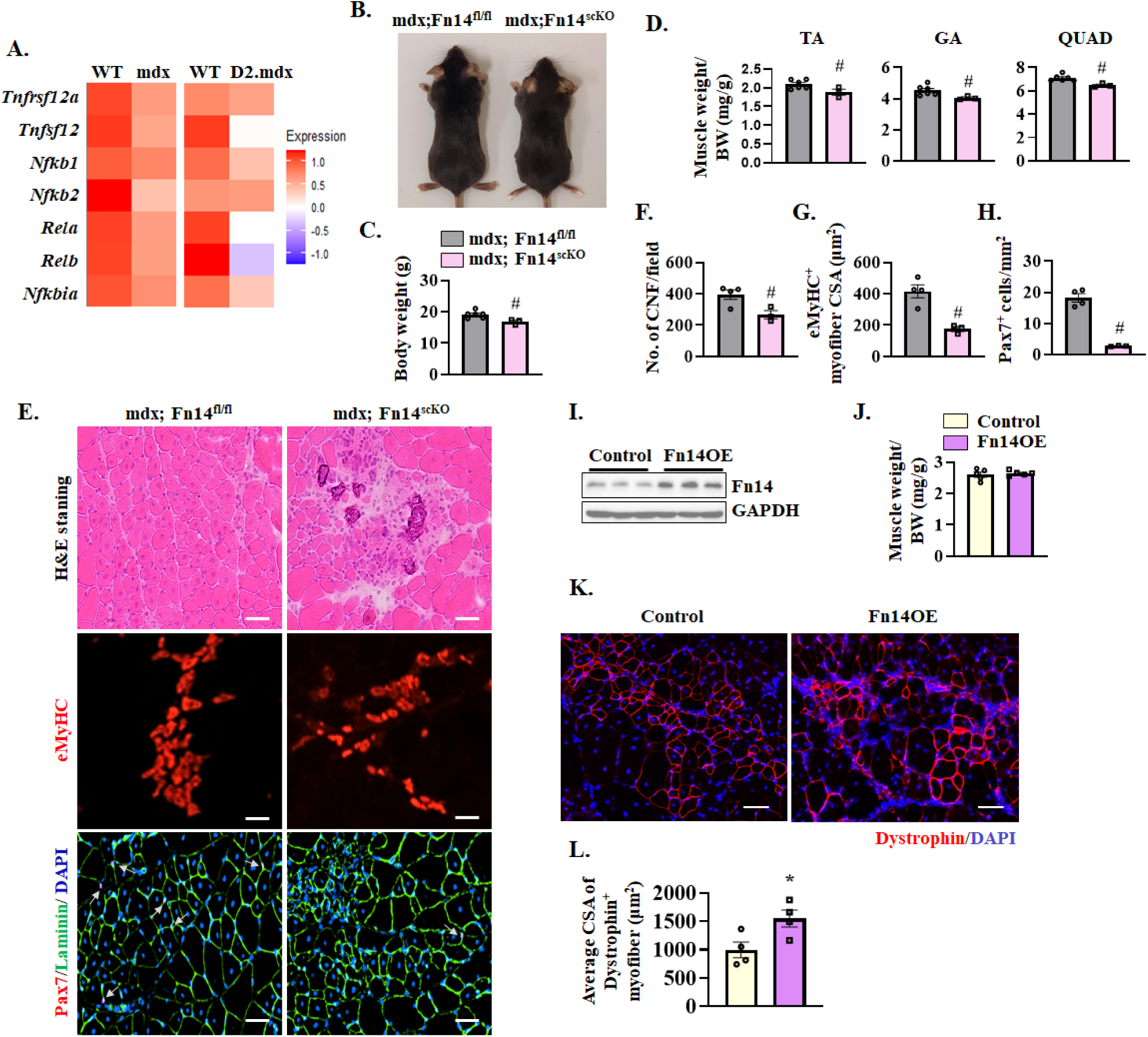
Satellite cell-specific ablation of Fn14 exacerbate dystrophic phenotype in mdx mice. **(A)** Heatmaps showing expression of Fn14 (gene name: *Tnfrsf12a*), TWEAK (gene name: *Tnfsf12*), *Nfkb1*, *Nfkb2*, *Rela*, *Relb*, and *Nfkbia* in muscle stem cells of WT and mdx, and WT and D2.mdx mice analyzed from a publicly available single cell dataset (GSE213925). **(B)** Representative pictures of 7-week-old mdx;Fn14^fl/fl^ and mdx;Fn14^scKO^ mice. Quantitative analysis of **(C)** body weight, and **(D)** wet weight of tibialis anterior (TA), gastrocnemius (GA), and quadriceps (QUAD) muscle normalized by BW of corresponding mice. **(E)** Representative photomicrographs of GA muscle sections of 7-week-old mdx;Fn14^fl/fl^ and mdx;Fn14^scKO^ mice after H&E staining, anti-eMyHC staining, or anti-Pax7/anti-laminin/DAPI staining. White arrows point to satellite cells in muscle sections. Scale bar: 50 µm. Quantification of **(F)** number of centronucleated fibers (CNF) per field, **(G)** average CSA of eMyHC^+^ myofibers, and **(H)** average number of Pax7^+^ cells per unit area. **(I)** Western blot demonstrating levels of Fn14 protein in myoblasts transduced with EGFP (control) or Fn14 overexpressing (Fn14OE) retrovirus. **(J)** TA muscle of 8-week-old mdx mice was injured by intramuscular injection of 1.2% BaCl_2_ solution. After 24 h, the muscle was injected with 1 × 10^6^ primary myoblasts stably transduced with control or Fn14OE retrovirus. After 28 days, the TA muscle was isolated and weighed. **(K)** Representative anti-dystrophin-stained images of TA muscle sections of mdx mice transplanted with control or Fn14OE myoblasts. **(L)** Quantitative analysis of average CSA of dystrophin^+^ myofibers in TA muscle of mdx mice transplanted with control or Fn14OE myoblasts. n= 3-6 mice in each group. All data are presented as mean ± SEM. #p ≤ 0.05, values significantly different from mdx;Fn14^fl/fl^ mice analyzed by unpaired Student *t* test. *p ≤ 0.05, values significantly different from TA muscle injected with control myoblasts by unpaired Student *t* test.

To further evaluate the role of Fn14 in satellite cell function, we performed myoblast transplantation and studied their engraftment in skeletal muscle of mdx mice (47, 48). Cultured myoblasts prepared from WT mice were stably transduced with control or Fn14-overexpressing (Fn14OE) retrovirus. Western blot analysis confirmed overexpression of Fn14 in myoblasts transduced with Fn14OE retrovirus (**Figure 8I**). Next, TA muscle of mdx mice was injured by intramuscular injection of 1.2% BaCl_2_ solution. After 24 h, the TA muscle was injected with control or Fn14-OE myoblasts. After 4 weeks, the TA muscle was isolated and analyzed. While there was no significant difference in wet weight of TA muscle injected with control or Fn14OE myoblasts (**Figure 8J**), there was a significant increase in the average CSA of dystrophin^+^ myofibers in the TA muscle injected with Fn14OE myoblasts compared with that injected with control myoblasts (**Figure 8, K and L**). Collectively, these results suggest that targeted ablation of Fn14 exacerbates myopathy in mdx mice potentially due to impairment in muscle regeneration and overexpression of Fn14 in exogenous myoblasts enhances their engraftment in dystrophic muscle of mdx mice.

## DISCUSSION

Skeletal muscle regeneration is a complex process that involves a number of intrinsic and extrinsic factors, which regulate the activation, proliferation, and differentiation of muscle progenitor cells (26). The Fn14 receptor is expressed on the cell surface of a variety of mesenchymal cells, including muscle progenitor cells (29, 31, 49). An earlier study showed that muscle regeneration is delayed in whole body Fn14-KO mice potentially due to reduced expression of chemokines, which promotes the recruitment of phagocytic cells after muscle damage (31). Consistently, we also recently reported that muscle regeneration is reduced in whole body Fn14-KO mice. Furthermore, we demonstrated that deletion of Fn14 in myoblasts (using Myod1-Cre line), but not in differentiated myofibers (using MCK-Cre line), inhibits skeletal muscle regeneration following acute injury and Fn14 regulates the myoblast fusion step during muscle repair (32). The role of Fn14 in myoblast fusion is also supported by another published report demonstrating that low doses of recombinant TWEAK protein promote fusion of cultured myoblasts (50).

Balance between self-renewal and differentiation is critical to simultaneously ensure maintenance of satellite cell pool while also generating differentiated progeny. Our experiments demonstrate that Fn14-mediated signaling is crucial for satellite cell homeostasis and function during skeletal muscle repair. Interestingly, we found that the levels of Fn14, but not TWEAK, increases in the satellite cells during muscle regeneration (**Figure 1**). Published reports suggest that while TWEAK is constitutively expressed and released in the circulation, the Fn14 receptor is expressed at a relatively low level in most cell types in uninjured healthy tissues. However, the activity of the TWEAK-Fn14 system is enhanced due to inducible local expression of Fn14 after injury in many tissues, which activates downstream signaling pathways regulating cell survival, proliferation, and differentiation (28–30). Therefore, it is likely that the increased expression of Fn14 in satellite cells is sufficient to activate the TWEAK-Fn14 signaling in satellite cells during muscle regeneration. Alternatively, it is also possible that elevated expression of Fn14 activates the downstream signaling pathways in satellite cells independently of TWEAK. In fact, a few studies have demonstrated that overexpression of Fn14 receptor, observed in certain cancers, can activate downstream signaling pathways even in the absence of TWEAK cytokine (29, 30).

Deletion of Fn14 in satellite cells of mice drastically inhibits regenerative myogenesis which is also evidenced by the findings that the expression of eMyHC protein persisted in Fn14^scKO^ mice even on day 21 post-injury (**Figure 2**, **Supplemental Figure 2**). Interestingly, the effect of satellite cell-specific deletion of Fn14 on muscle regeneration observed in this study (**Figure 2**) is much more pronounced than the whole body Fn14-KO mice which also shows delay in muscle regeneration after acute injury (31, 32). These differences in the severity of regeneration defects between whole body and conditional satellite cell-specific Fn14 knockout mice could be attributed to the fact that in whole body Fn14-KO mice, the deficiency of Fn14 in satellite cells may be partially compensated by other factors present during embryonic development.

Moreover, it is possible that the lack of Fn14 signaling in other cell types also influences injured muscle microenvironment, including inflammatory response, which may compensate for the lack of Fn14 in satellite cells to some extent. For our present study, we used tamoxifen-inducible satellite cell-specific Fn14-knockout mice, which allows understanding the role of satellite cell Fn14 signaling in skeletal muscle regeneration that is devoid of any confounding effects of Fn14 deficiency during muscle development and in other cell types that are present in regenerating skeletal muscle.

Our results demonstrate that Fn14 plays a crucial role in the self-renewal and proliferation of satellite cells following injury-induced muscle regeneration. There was a dramatic reduction in the abundance of satellite cells in regenerating muscle of Fn14^scKO^ mice (**Figure 3**). Furthermore, double injury to TA muscle also led to enhanced deficits in muscle regeneration and reduction in the number of satellite cells, suggesting that Fn14 is critical for replenishing the pool of satellite cells after injury (**Figure 5**). The effects of Fn14 deletion in satellite cell proliferation are also evidenced by the *ex vivo* and *in vitro* findings. Several signaling pathways have now been identified that regulate satellite cell fate and their ability to repair injured myofibers. Notch signaling is essential for the self-renewal and proliferation of satellite cells and to inhibit their differentiation into myogenic lineage through repressing the levels of MyoD (10, 11, 13, 14).

Moreover, Notch3 is required for maintaining the quiescent satellite cell population in adult skeletal muscle (51). Genetic ablation of either Notch1 or Notch2 alone in activated satellite cells attenuates population expansion and self-renewal but induces premature myogenesis (15).

Intriguingly, our results demonstrate that the levels of Pax7 and cleaved Notch1 protein are reduced in cultured Fn14-KO myoblasts. Furthermore, the gene expression of various components of Notch signaling, including receptors, ligands, and target genes are significantly reduced in regenerating muscle of Fn14^scKO^ mice compared to corresponding control mice (**Figure 5, G-I**) suggesting that Fn14-mediated signaling crosstalk with Notch signaling to promote the self-renewal and proliferation of satellite cells in adult skeletal muscle.

A striking observation of the present study was that the gene expression of multiple components of the JAK-STAT signaling pathway, including ligands (e.g., IL-6) and target genes was significantly up-regulated whereas the expression of negative regulators (e.g., Socs2, Socs3, and Socs5) of this pathway was reduced in Fn14-KO myoblast cultures compared to corresponding controls (**Figure 7A**). Our experiments further demonstrate that satellite cell-specific deletion of Fn14 leads to the activation of STAT3 signaling in regenerating muscle as well as in cultured myoblasts (**Figure 7, B and C**). It has been previously reported that extracellular molecules, such as IL-6 and leukemia inhibitory factor, activate STAT3 which augments myogenic differentiation through interaction with MyoD transcription factor (52). Recent studies have demonstrated that genetic ablation of STAT3 in satellite cells augments their expansion during muscle regeneration but inhibits their differentiation (22). The study showed that transient Stat3 inhibition promotes satellite cell expansion and improves tissue repair in both aged and dystrophic muscle of mice (22). Similarly, another study demonstrated that genetic knockdown or pharmacological inhibition of Jak2 and Stat3 stimulates the proliferation of satellite cells and their engraftment *in vivo* (23). In contrast, Zhu et al demonstrated that physiological levels of Stat3 are essential for the self-renewal of satellite cells during muscle regeneration (24). Our results suggest that aberrant activation of Stat3 is responsible for the reduced proliferation and precocious differentiation of Fn14-deficient satellite cells (**Figure 7**). Interestingly, we also observed that knockdown of Stat3 augments the expression of Fn14 in cultured myoblasts (**Figure 7F**) suggesting the presence of a regulatory circuitry between Fn14 and Stat3 in the regulation of satellite cell expansion and differentiation. Altogether, our experiments demonstrate that Fn14 regulates both Notch and STAT3 signaling, which may be important mechanisms regulating the satellite cell content and function during regenerative myogenesis.

Future studies should investigate the molecular interactions through which Fn14 regulates these pathways in satellite cells.

The critical role of Fn14 receptor-mediated signaling in satellite cell function is further supported by our findings in the mdx mice. DMD is a genetic muscle disorder, which involves chronic degeneration and regeneration of myofibers and eventually replacement of muscle tissue by connective tissues (53–55). The mdx mouse model is widely used to study muscle regeneration in disease conditions and in response to repeated cycles of injury (56–58). Targeted deletion of Fn14 in mdx mice exacerbates myopathy potentially due to a reduced number of satellite cells, which results in the impairment of regeneration of injured myofibers (**Figure 8**). Interestingly, we also found that the gene expression of Fn14 and its downstream targets is significantly reduced in muscle progenitor cells of both mdx and clinically relevant D2.mdx models of DMD suggesting that Fn14 is involved in the etiology of DMD (**Figure 8A**). The beneficial role of Fn14-mediated signaling in satellite cell function is supported by our findings that overexpression of Fn14 in exogenous muscle progenitor cells resulted in their enhanced engraftment when transplanted into the dystrophic muscle of mdx mice (**Figure 8, I-L**). Since Fn14 also promotes myoblast fusion through augmenting the gene expression of various profusion molecules (32), it is possible that in addition to improving abundance and myogenic potential of satellite cells, overexpression of Fn14 in muscle progenitor cells also augments their fusion to injured myofibers of dystrophic muscle resulting in a better engraftment.

The results of our present study and previously published reports demonstrate that Fn14 signaling in muscle progenitor cells is required for their proliferation and fusion with injured myofibers (32, 50). It is notable that chronic stimulation of Fn14 receptor by its ligand TWEAK has been shown to inhibit satellite cell self-renewal and myotube formation in cultured myoblasts after induction of differentiation (31, 59–61). Chronic treatment with high levels of TWEAK protein might have broad non-specific effects leading to impairment in myogenesis and muscle atrophy (30, 59, 60, 62, 63). By contrast, overexpression of Fn14 in myoblasts, even in the absence of TWEAK cytokine, is sufficient to activate downstream signaling and augment myoblast fusion without affecting myogenic differentiation (32) suggesting that overexpression of Fn14 is a valid approach to improve satellite cell myogenic potential and their fusion with injured myofibers to improve muscle repair.

For this study, we used only male mice because in our initial studies we observed similar deficits in muscle regeneration in both male and female Fn14^scKO^ mice. However, there are published reports suggesting sex-specific differences in muscle function, regeneration, and metabolism (64, 65). Further experimentation is needed to determine whether there are gender differences regarding the role of Fn14 in regulating satellite cell function and various aspects of muscle regeneration and function. Our studies were performed with TA and EDL muscles, which predominately contain fast-twitch myofibers. Future studies will investigate whether Fn14-mediated signaling in satellite cells similarly regulates regeneration in the soleus and gastrocnemius muscles, which contain both slow-and fast-twitch myofibers.

In summary, the present study has identified a previously unrecognized role of Fn14 in regulating satellite cell homeostasis and function. Fn14 is necessary for maintaining the satellite cell pool, sustaining the myogenic potential upon stimulation, and regulating self-renewal following activation. While more investigations are needed, our study suggests that augmenting the levels of Fn14 in satellite cells could be an important therapeutic approach for various muscle wasting conditions, such as aging and various degenerative muscle disorders.

## MATERIALS AND METHODS

### Animals

C57BL/6 and mdx mice were purchased from Jackson Labs. Whole-body Fn14- knockout (i.e., Fn14-KO) mice have been previously described (31). Floxed Fn14 (i.e., Fn14^fl/fl^) mice were generated by Taconic by inserting loxP sites upstream of exon 2 and downstream of exon 4 of Fn14 as described (66). Floxed Fn14 mice were crossed with *Pax7-CreERT2* mice (Jax strain: B6;129-Pax7^tm2.1(cre/ERT2)Fan^/J) to generate satellite cell-specific Fn14-knockout (Fn14^scKO^) mice. Similarly, mdx mice were crossed with Fn14^scKO^ to generate mdx;Fn14^scKO^ mice and littermate mdx; Fn14^fl/fl^ mice. All mice were in the C57BL/6 background. Mice were housed in an environmentally controlled room (23 °C, 12-h light-dark cycle) with ad libitum access to food and water. Genotype of mice was determined by performing PCR from tail DNA. We used 7-8 weeks old male mice for our experimentation. For Cre-mediated inducible deletion of Fn14 in satellite cells, Fn14^scKO^ mice were given intraperitoneal (i.p.) injection of tamoxifen (10 mg per Kg body weight) in corn oil for four consecutive days and kept on tamoxifen-containing standard chow (Harlan Laboratories,Madison, WI) for the entire duration of the experiment. For the mdx;Fn14^scKO^ mice, injections of tamoxifen were given at the age of 3 weeks, for four days to induce Cre-mediated recombination and mice were kept on a tamoxifen-containing chow for the entire duration of the study and analyzed at the age of 7 weeks. Fn14^fl/fl^ and mdx;Fn14^fl/fl^ mice were also injected with tamoxifen following the same regimen and served as controls. All surgeries were performed under anesthesia, and every effort was made to minimize suffering.

### Skeletal muscle injury

TA muscle of adult mice was injected with 50 µl of 1.2% BaCl_2_ (Sigma Chemical Co.) dissolved in saline to induce necrotic injury. The mice were euthanized at different time points after injury and TA muscle was isolated for biochemical and morphometric analysis.

### Histology and morphometric analysis

Uninjured or injured TA muscle was isolated from mice and sectioned with a microtome cryostat. For the assessment of muscle morphology and quantification of myofiber cross-sectional area (CSA), 8-μm-thick transverse sections of TA muscle were stained with Hematoxylin and Eosin (H&E). Sirius red (StatLab) staining was performed to visualize fibrosis. All stained and immunofluorescence labeled sections were visualized, and images were captured using an inverted microscope (Nikon Eclipse Ti-2E Inverted Microscope), a digital camera (Digital Sight DS-Fi3, Nikon) and NIS Elements AR software (Nikon) at room temperature. The NIH ImageJ software was used for quantitative analysis and image levels were equally adjusted using Photoshop CS6 software (Adobe).

### Immunohistochemistry

For immunohistochemistry studies, frozen TA muscle sections were fixed in acetone or 4% paraformaldehyde (PFA) in PBS, blocked in 2% bovine serum albumin in PBS for 1 h, followed by incubation with anti-Pax7 or anti-eMyHC and anti-laminin in blocking solution at 4 °C overnight under humidified conditions. The sections were washed briefly with PBS before incubation with goat anti-mouse Alexa Fluor 594, or 568 and goat anti-rabbit Alexa Fluor 488 secondary antibody for 1 h at room temperature and then washed three times for 15 min with PBS. Nuclei were counterstained with DAPI. The slides were mounted using fluorescence medium (Vector Laboratories).

### Isolation and culturing of primary myoblasts

Primary myoblasts were prepared from the hind limbs of 8-week-old male mice following a protocol as described (67, 68). Cultures were transfected with control or STAT3 siRNA (SantaCruz Biotechnology) using Lipofectamine RNAiMAX Transfection Reagent (ThermoFisher Scientific).

### Generation and use of retroviruses

A pBABE-Puro empty vector and a pBABE-Puro-EGFP plasmid were purchased from Addgene. Mouse Fn14 cDNA was isolated and ligated at BamHI and SalI sites in the pBABE-Puro plasmid. The integrity of the cDNA was confirmed by performing DNA sequencing. About 5 × 10^6^ Platinum-E packaging cells (Cell Biolabs, Inc.) were transfected with 10 μg of pBABE-Puro-Fn14 or pBABE-Puro-EGFP using FuGENE-HD (Promega). After 24 h of transfection, the medium was replaced with growth media supplemented with 10% FBS. After 48 h of transfection, viral supernatants were collected, filtered through 0.45-micron filters, and added to primary myoblasts in growth media containing 10 μg/ml polybrene. After two successive retroviral infections, cells were selected using 1.2 μg/ml puromycin for 1 week.

### Myoblast transplantation

TA muscle of 8-week-old mdx mice was injured by an intramuscular injection of 50 μl 1.2% BaCl_2_ solution. After 24 h, the muscle was injected with 1 × 10^6^ primary myoblasts prepared from WT mice and transduced with the retrovirus expressing EGFP (control) or Fn14 cDNA. Twenty-eight days after transplantation, the TA muscle was isolated and transverse sections made were immunostained for dystrophin protein, followed by measuring average CSA of dystrophin^+^ myofibers (48).

### Isolation, culture, and staining of single myofibers

Single myofibers were isolated from EDL muscle after digestion with collagenase II (Sigma-Aldrich) and trituration as previously described (69). Suspended fibers were cultured in 60-mm horse serum–coated plates in DMEM supplemented with 10% FBS (Invitrogen), 2% chicken embryo extract (Accurate Chemical and Scientific Corporation), 10 ng/ml basis fibroblast growth factor (PeproTech), and 1% penicillin-streptomycin for 3 days. Freshly isolated fibers and cultured fibers were fixed in 4% PFA and stained with anti-Pax7 (1:10, Developmental Studies Hybridoma Bank [DSHB]) and MyoD (1:200, sc-377460, Santa Cruz Biotechnology Inc.).

### Cell proliferation assay

Satellite cell or myoblast proliferation was assayed by labeling the cells with EdU for 60 min using Click-iT EdU Cell Proliferation Assay kit (Invitrogen) as previously described (61). To study proliferation of muscle progenitor cells *in vivo*, the mice were given an intraperitoneal injection of EdU (4 μg per gram body weight) at day 3 post BaCl2-medited injury of TA muscle. On day 11 after EdU injection, the mice were euthanized, and TA muscle was isolated and transverse sections were made. The TA muscle sections were subsequently immunostained with anti-Laminin for marking boundaries of myofibers and processed for the detection of EdU^+^ nuclei. In case of single myofiber or myoblast cultures, EdU pulse-labelling was performed for 60 min. The EdU^+^ nuclei on muscle sections were detected as instructed in the Click-iT EdU Alexa Fluor 488 Imaging Kit (Invitrogen). DAPI stain was used to identify nuclei. Finally, images were captured and the number of intramyofiber EdU^+^ myonuclei per myofiber, percentage of 2 or more EdU^+^ centrally nucleated fibers, and percentage of EdU^+^ myonuclei/total DAPI^+^ nuclei were evaluated using NIH ImageJ software.

Three to four different sections from mid-belly of each muscle were included for analysis.

### Fluorescence-activated cell sorting (FACS)

Satellite cells from injured TA muscle were analyzed by FACS. After muscle digestion with collagenase type II, cells were incubated in DMEM (supplemented with 2% FBS). For satellite cell quantification or isolation from heterogeneous cell population, cells were immunostained with antibodies against CD45, CD31, Sca-1, and Ter-119 for negative selection (all PE conjugated), and with α7-integrin (APC conjugated) for positive selection. To detect Fn14 expression in satellite cells, the cells were also incubated with anti-Fn14 and then incubated with anti-mouse IgG2b FITC secondary antibody. FACS analysis was performed on a C6 Accuri cytometer (BD Biosciences) equipped with 3 lasers. The output data was processed, and plots were prepared using Flow Jo software (BD Biosciences).

### Satellite cell isolation

Satellite cells were isolated from uninjured and injured TA muscles of WT, Fn14^fl/fl^ and Fn14^scKO^ mice using Satellite Cell Isolation Kit, mouse (Miltenyi Biotec).

Briefly, TA muscle of 8-week-old mice was injured by an intramuscular injection of 50 μl 1.2% BaCl_2_ solution. After 5 or 21 days, muscles were isolated, washed in PBS, minced into coarse slurry and enzymatically digested at 37°C for 1 h by adding 2% collagenase II (Gibco, Life Technologies). The digested slurry was spun, pelleted and triturated several times and then passed through a 70 µm and then 30 µm cell strainer (BD Falcon). The filtrate was spun at 1,000g and the cell pellets were used for satellite cell isolation using the manufacturer’s protocol (Miltenyi Biotec).

### Lactate dehydrogenase (LDH) assay

The amount of LDH in culture supernatants was measured using a commercially available LDH Cytotoxicity Assay kit following the protocol suggested by the manufacturer (Thermo Scientific Life Sciences).

### RNA isolation and qRT-PCR

RNA isolation and qRT-PCR were performed following a standard protocol as described (67). In brief, total RNA was extracted from injured TA muscle or freshly isolated satellite cells of mice using TRIzol reagent (Thermo Fisher Scientific) and RNeasy Mini Kit (Qiagen, Valencia, CA, USA) according to the manufacturers’ protocols. First-strand cDNA for PCR analyses was made with a commercially available kit (iScript cDNA Synthesis Kit, Bio-Rad Laboratories). The quantification of mRNA expression was performed using the SYBR Green dye (Bio-Rad SsoAdvanced - Universal SYBR Green Supermix) method on a sequence detection system (CFX384 Touch Real-Time PCR Detection System - Bio-Rad Laboratories). The sequence of the primers is described in **Supplemental Table 1**. Data normalization was accomplished with the endogenous control (β-actin), and the normalized values were subjected to a 2^-ΔΔCt^ formula to calculate the fold change between control and experimental groups.

### Western Blot

TA muscle of mice or cultured primary myoblasts were washed with PBS and homogenized in lysis buffer (50 mM Tris-Cl (pH 8.0), 200 mM NaCl, 50 mM NaF, 1 mM dithiothreitol, 1 mM sodium orthovanadate, 0.3% IGEPAL, and protease inhibitors).

Approximately, 100 μg protein was resolved on each lane on 8-12% SDS-PAGE gel, transferred onto a nitrocellulose membrane, and probed using specific primary antibody (**Supplemental Table 2**). Bound antibodies were detected by secondary antibodies conjugated to horseradish peroxidase (Cell Signaling Technology). Signal detection was performed by an enhanced chemiluminescence detection reagent (Bio-Rad). Approximate molecular masses were determined by comparison with the migration of prestained protein standards (Bio-Rad).

### Analysis of single-cell RNA sequencing dataset

The publicly available dataset (GSE143437) raw data was processed using Seurat package on R software (v4.2.2) to analyse gene expression of specific molecules in satellite cells of wild-type mice at different time points after injury.

Similarly, another publicly available dataset (GSE213925) was analyzed to study the gene expression of the same molecules in satellite cells of WT, mdx, and D2.Mdx mice.

### RNA sequencing and data analysis

Total RNA from control, WT and Fn14-KO cultures was extracted using TRIzol reagent (Thermo Fisher Scientific) using the RNeasy Mini Kit (Qiagen, Valencia, CA, USA) according to the manufacturers’ protocols. The mRNA-seq library was prepared using poly (A)-tailed enriched mRNA at the UT Cancer Genomics Center using the KAPA mRNA HyperPrep Kit protocol (KK8581, Roche, Holding AG, Switzerland) and KAPA Unique Dual-indexed Adapter kit (KK8727, Roche). The Illumina NextSeq550 was used to produce 75 base paired end mRNA-seq data at an average read depth of ∼38M reads/sample.

RNA-seq fastq data was processed using CLC Genomics Workbench 20 (Qiagen), followed by trimming of adapter sequences and alignment to the mouse reference genome Refseq GRCm39.105 from the Biomedical Genomics Analysis Plugin 20.0.1 (Qiagen). Normalization of RNA-seq data was performed using trimmed mean of M-values. Genes with fold change (FC) ≥ 1.5 (or log2FC ≥ 0.5) and p-value <0.05 were assigned as differentially expressed genes (DEGs). We have previously published results based on the dataset (accession code mentioned in the data availability section) (32). We utilized a subset of these samples (only 0 h time point) to evaluate gene expression of differente molecules associated with satellite cell function.

Heatmaps were generated by using heatmap.2 function (Gu & Hubschmann, 2022) using z-scores calculated based on transcripts per million (TPM) values. Average of absolute TPM values for control group are provided in **Supplemental Table 3**. TPM values were converted to log (TPM+1) to handle zero values. Genes involved in specific pathways were manually selected for heatmap expression plots.

### Statistical analyses and experimental design

The sample size was calculated using power analysis methods for a priori determination based on the s.d. and the effect size was previously obtained using the experimental procedures employed in the study. For animal studies, we calculated the minimal sample size for each group as eight animals. Considering a likely drop-off effect of 10%, we set sample size of each group of six mice. For some experiments, three to four animals were sufficient to obtain Statistically significant differences. Animals with same sex and same age were employed to minimize physiological variability and to reduce s.d. from mean.

The exclusion criteria for animals were established in consultation with a veterinarian and experimental outcomes. In case of death, skin injury, ulceration, sickness, or weight loss of > 10%, the animal was excluded from analysis. Tissue samples were excluded in cases such as freeze artifacts on histological sections or failure in extraction of RNA or protein of suitable quality and quantity. We included animals from different breeding cages by random allocation to the different experimental groups. All animal experiments were blinded using number codes until the final data analyses were performed. Statistical tests were used as described in the Figure legends. Results are expressed as mean ± SEM. Statistical analyses used two-tailed Student’s t-test or 2-way ANOVA followed by Tukey’s multiple-comparison test to compare quantitative data populations with normal distribution and equal variance. A value of p ≤ 0.05 was considered statistically significant unless otherwise specified.

## Study approval

All animal procedures were conducted in strict accordance with the institutional guidelines and were approved by the Institutional Animal Care and Use Committee and Institutional Biosafety Committee of the University of Houston (PROTO201900043).

## Data availability

All relevant data related to this manuscript are available from the authors upon reasonable request. All the raw data files for RNA-seq experiment can be found on NCBI SRA repository using the accession code <u>PRJNA999372</u>.

## Acknowledgements

We are greatful to Dr. Linda Burkly of Biogen Inc. (Cambridge, MA) for providing floxed Fn14 and whole body Fn14-knockout mice.This work was supported by National Institute of Health grant AR081487 to AK. We thank the technical support from the Cancer Prevention and Research Institute of Texas (CPRIT RP180734) core for RNA-seq experiment.

## Authors’ contribution

A.K. designed the work. M.T.S. and A.S.J. performed all the experiments. M.T.S. wrote the first draft of the manuscript and A.K. edited and finalized the manuscript.

## Conflict of Interest

The authors declare no competing interests.

## Notes

https://www.ncbi.nlm.nih.gov/bioproject/PRJNA999372

### Competing Interest Statement

The authors have declared no competing interest.

### Summary of Updates

New data have been added in Figure 1, Figure 4, Figure 6, Figure S1, and Figure S3. The text has been modified in the results section accordingly. Also the Discussion section of the manuscript has also been updated.

