## Supplemental Figures, 1-5 and Supplemental Table 1-3. for "Fibroblast growth factor-inducible 14 regulates satellite cell self-renewal and expansion during skeletal muscle repair"

### <u>Supplemental Data File</u>

**By**

**Meiricris Tomaz da Silva, Aniket S. Joshi, and Ashok Kumar**

This file contains Supplemental Figures, 1-5 and Supplemental Table 1-3.

### Supplemental Figures' and Legends

**Supplemental Figure 1.**

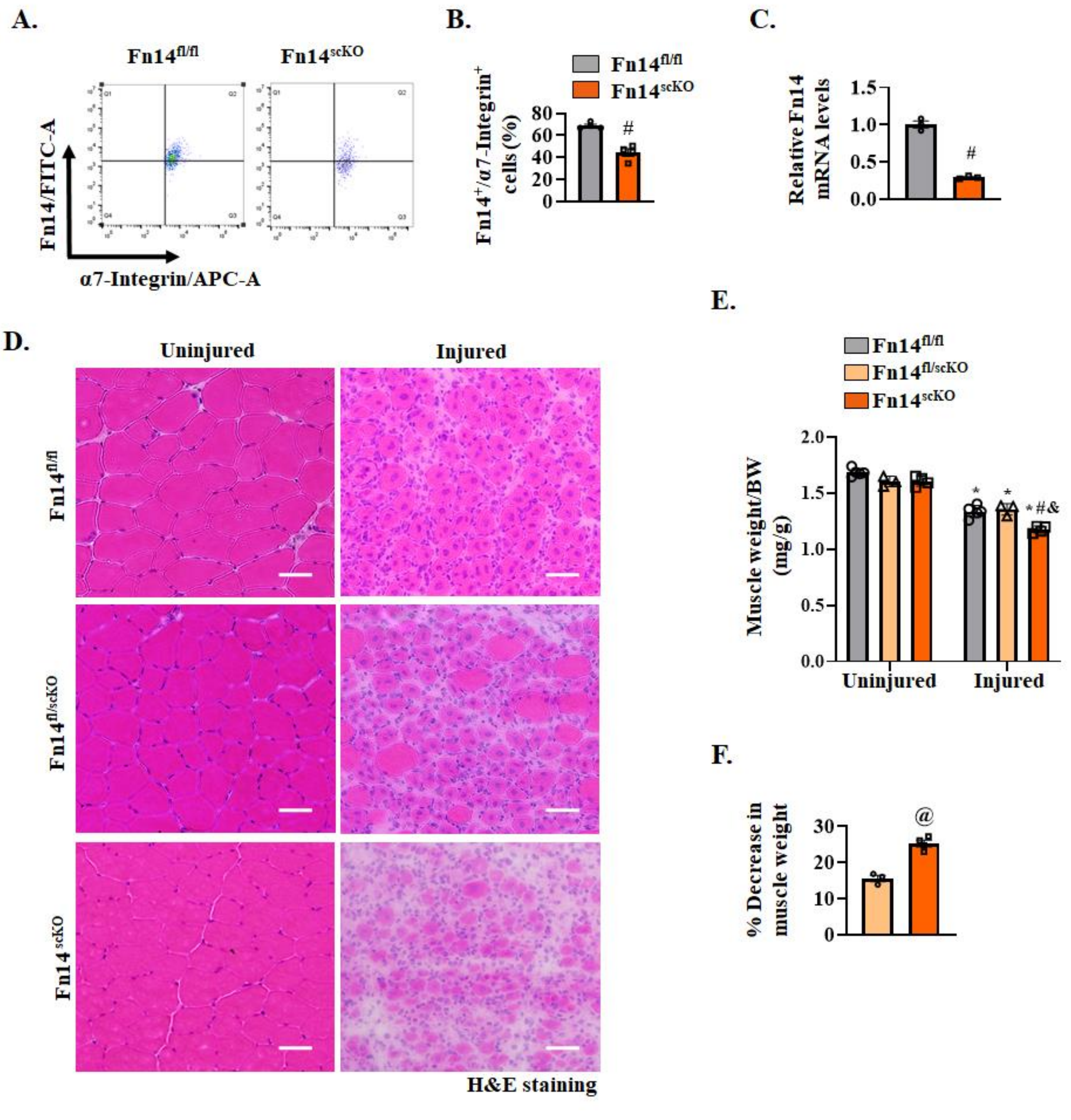


**Supplemental FIGURE 1**. **Fn14 signaling in satellite cells mediates muscle regeneration.** **(A)** Primary mononuclear cells were isolated from TA muscle of Fn14$^{fl/fl}$ and Fn14$^{scKO}$ mice and subjected to FACS analysis for the expression of α7-integrin and Fn14. Representative scatter plots of FACS-based demonstrating Fn14$^{+}$ and α7-integrin$^{+}$ cells in 5d-injured TA muscle of Fn14$^{fl/fl}$ and Fn14$^{scKO}$ mice. **(B)** Quantification of the percentage of Fn14$^{+}$/α7-integrin$^{+}$ cells in 5d-injured TA muscle of Fn14$^{fl/fl}$ and Fn14$^{scKO}$ mice assayed by FACS. **(C)** Relative mRNA levels of Fn14 in isolated satellite cells of 5d-injured TA muscle of $Fn14^{fl/fl}$ and $Fn14^{scKO}$ mice. n=3-4 mice in each group. #$p \leq 0.05$, values significantly different from corresponding 5d-injured TA muscle of $Fn14^{fl/fl}$ mice analyzed by unpaired Student *t* test. **(D)** Representative photomicrographs of H&E stained sections of uninjured and injured TA muscles of $Fn14^{fl/fl}$, $Fn14^{fl/wt}$; Pax7-CreERT2 ($Fn14^{fl/scKO}$) and $Fn14^{scKO}$ mice. Scale bar, 50 µm. **(E)** Uninjured and 5d-injured TA muscle wet weight normalized by body weight (BW) of $Fn14^{fl/scKO}$ and $Fn14^{scKO}$. **(F)** Percentage decrease in wet weight of TA muscle of $Fn14^{fl/scKO}$ and $Fn14^{scKO}$ after injury. n= 3-4 mice in each group. All data are presented as mean ± SEM. *$p \leq 0.05$, values significantly different from contralateral uninjured muscle of $Fn14^{fl/fl}$ or $Fn14^{fl/scKO}$ mice; $^{\&}p \leq 0.05$, values significantly different from corresponding 5d-injured TA muscle of $Fn14^{fl/fl}$ mice analyzed by two-way ANOVA followed by Tukey's multiple comparison test. $^{@}p \leq 0.05$, values significantly different from corresponding $Fn14^{fl/scKO}$ mice analyzed by unpaired Student *t* test.

**Supplemental Figure 2.**

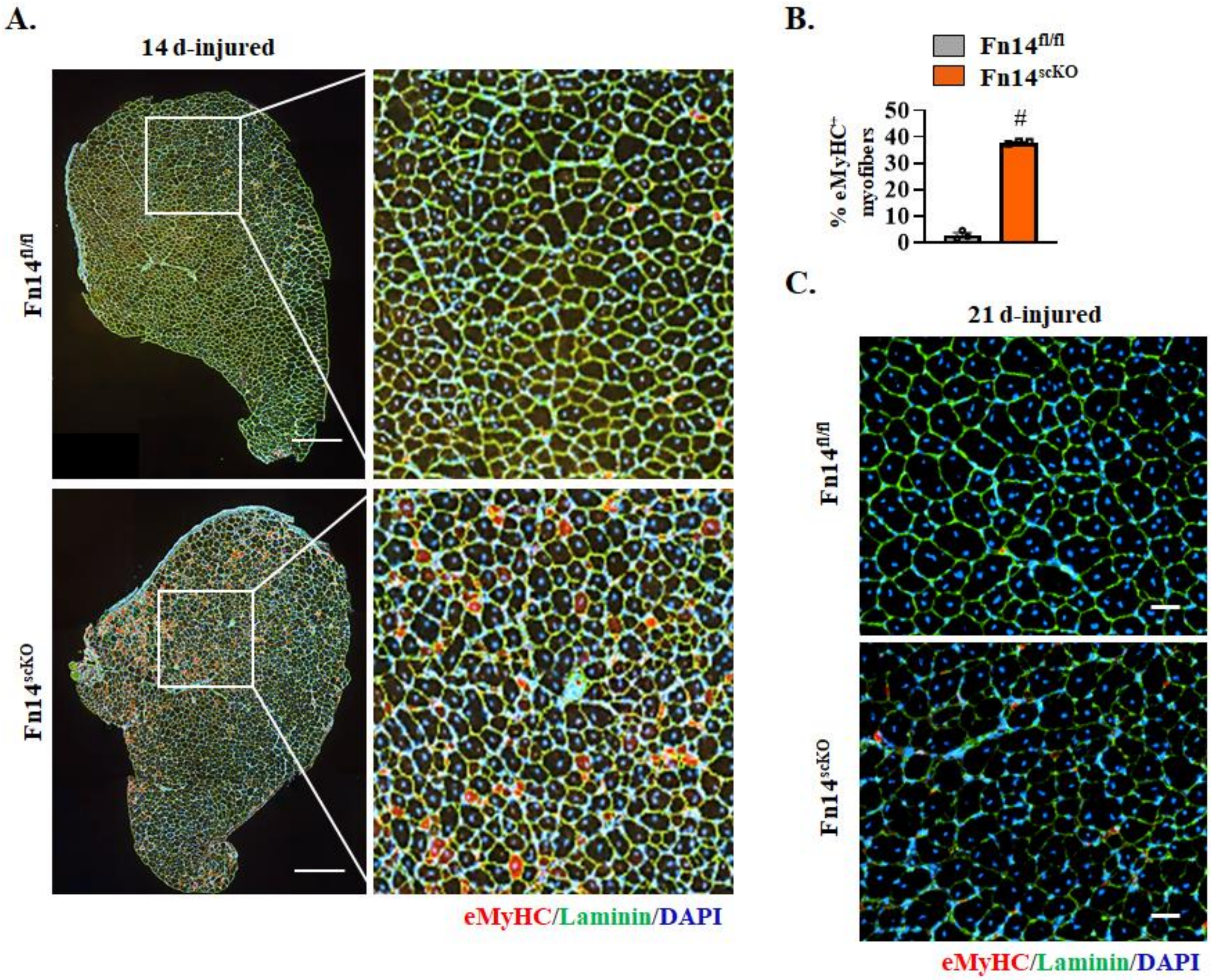


**Supplemental FIGURE 2**. **Deletion of Fn14 in satellite cells delays muscle regeneration. (A)** Representative photomicrographs of transverse sections of 14d-injured TA muscle of $Fn14^{fl/fl}$ and $Fn14^{scKO}$ mice after immunostaining for eMyHC and laminin protein and DAPI staining. Left panel: whole muscle section; right panel: magnified view of the selected region. Scale bar: 400 µm. **(B)** Quantification of percentage of $eMyHC^+$ myofibers in 14d-injured TA muscle section of $Fn14^{fl/fl}$ and $Fn14^{scKO}$ mice. **(C)** Representative photomicrographs of transverse sections of 21d-injured TA muscle of $Fn14^{fl/fl}$ and $Fn14^{scKO}$ mice after immunostaining for eMyHC and laminin protein and DAPI staining. Scale bar: 50 µm. n= 3 mice in each group. Data are presented as mean ± SEM. #p ≤ 0.05, values significantly different from corresponding muscle of $Fn14^{fl/fl}$ mice analyzed by unpaired Student *t* test.

**Supplemental Figure 3.**

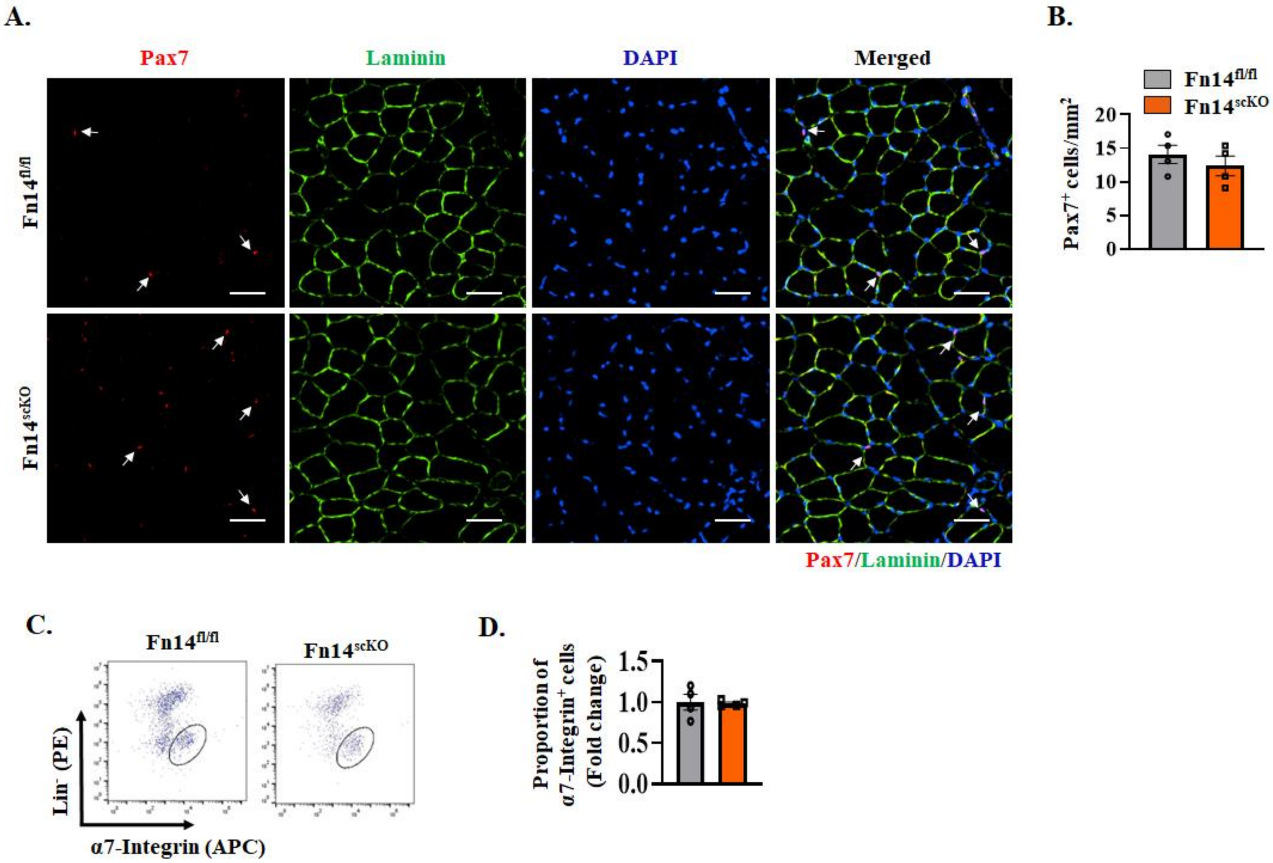


**Supplemental FIGURE 3**. **Targeted deletion of Fn14 does not affect satellite cell number in uninjured muscle. (A)** Representative photomicrographs of uninjured TA muscle sections of Fn14$^{fl/fl}$ and Fn14$^{scKO}$ mice after immunostaining for Pax7 (red) and laminin (green) protein. Nuclei were counterstained with DAPI. Scale bar: 50 µm. White arrows point to Pax7$^{+}$ satellite cells. **(B)** Average number of Pax7$^{+}$ cells per unit area in uninjured TA muscle of Fn14$^{fl/fl}$ and Fn14$^{scKO}$ mice. n= 4 mice in each group. **(C)** Single cell suspensions were isolated from TA muscle of Fn14$^{fl/fl}$ and Fn14$^{scKO}$ mice were subjected to FACS analysis for satellite cells. Representative FACS dot plots demonstrating the percentage of α7-integrin$^{+}$ cells in uninjured TA muscle of Fn14$^{fl/fl}$ and Fn14$^{scKO}$ mice. **(D)** Quantification of α7-integrin$^{+}$ satellite cells in uninjured TA muscle of Fn14$^{fl/fl}$ and Fn14$^{scKO}$ mice assayed by FACS. All data are presented as mean ± SEM. No statistically significant difference was obtained between the two genotypes by unpaired Student *t* test.

**Supplemental Figure 4.**

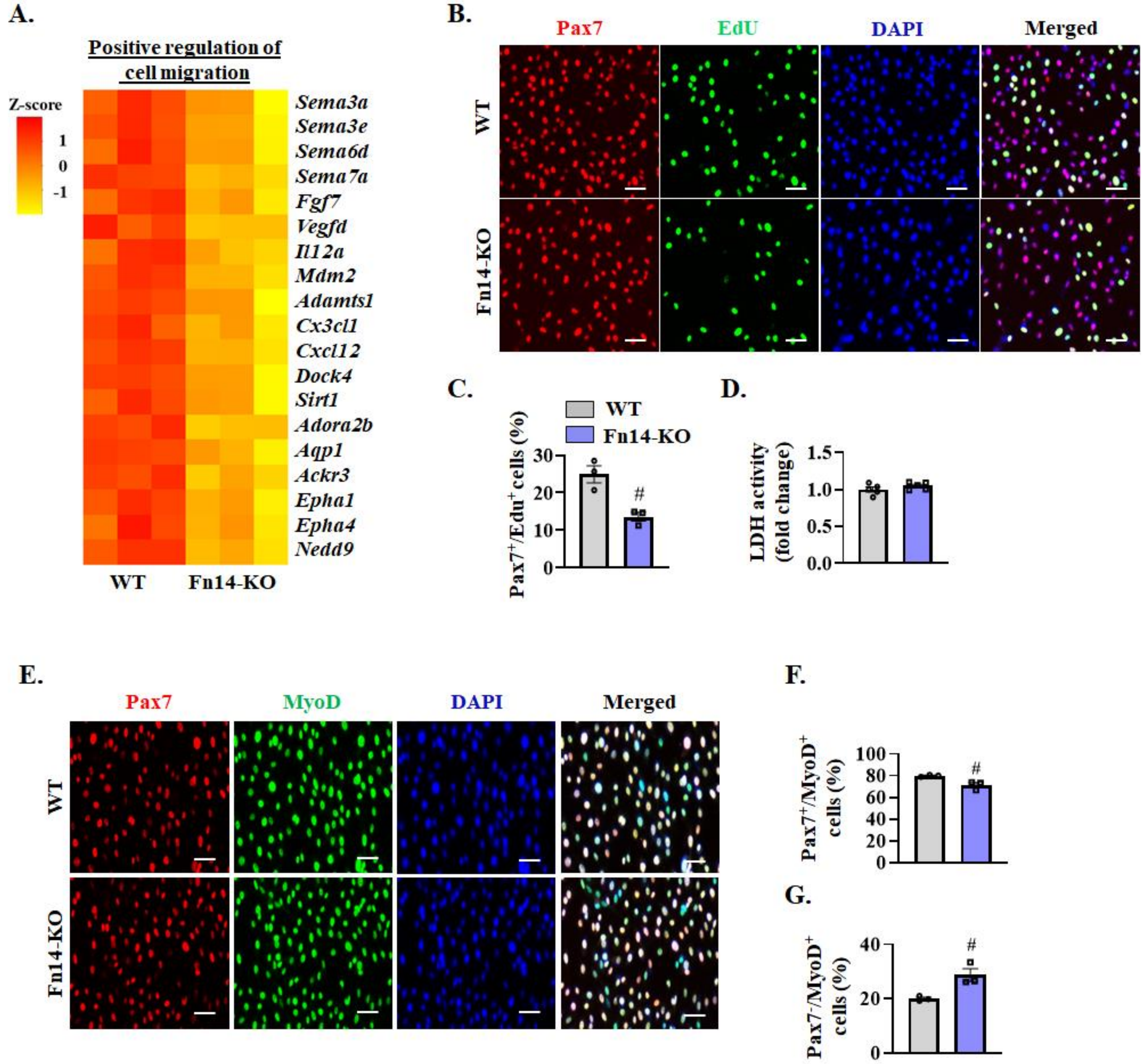


**Supplemental FIGURE 4**. **Effect of deletion of Fn14 on satellite cell survival and proliferation. (A)** Heatmap representing selected genes associated with the positive regulation of cell migration in WT and Fn14-KO cultures generated by analysis of RNA-seq dataset. **(B)** WT and Fn14-KO myoblasts were seeded at equal number and pulse-labeled with EdU for 60 min followed detection of $EdU^+$ nuclei and immunostaining for Pax7 protein. Representative photomicrographs of merged images are presented here. Scale bar: 50 µm. **(C)** Quantification of percentage of $Pax7^+$/$EdU^+$ cells in WT and Fn14-KO cultures. **(D)** Relative amounts of lactate dehydrogenase (LDH) in supernatants of WT and Fn14-KO cultures.  n= 3 biological replicates in each group. **(E)** Representative photomicrographs of WT and Fn14-KO primary myoblast cultures after immunostaining for Pax7 and MyoD protein. Nuclei were counterstained by DAPI. Scale bar: 50 µm. Quantification of percentage of **(F)** $Pax7^+$/$MyoD^+$ (proliferating), and **(G)** $Pax7^-$/$MyoD^+$ (differentiating) cells in WT and Fn14-KO cultures. n= 3 biological replicates in each group. All data are presented as mean ± SEM. #$p \leq 0.05$, values significantly different from WT cultures analyzed by unpaired Student *t* test.

#### Supplemental Figure 5.

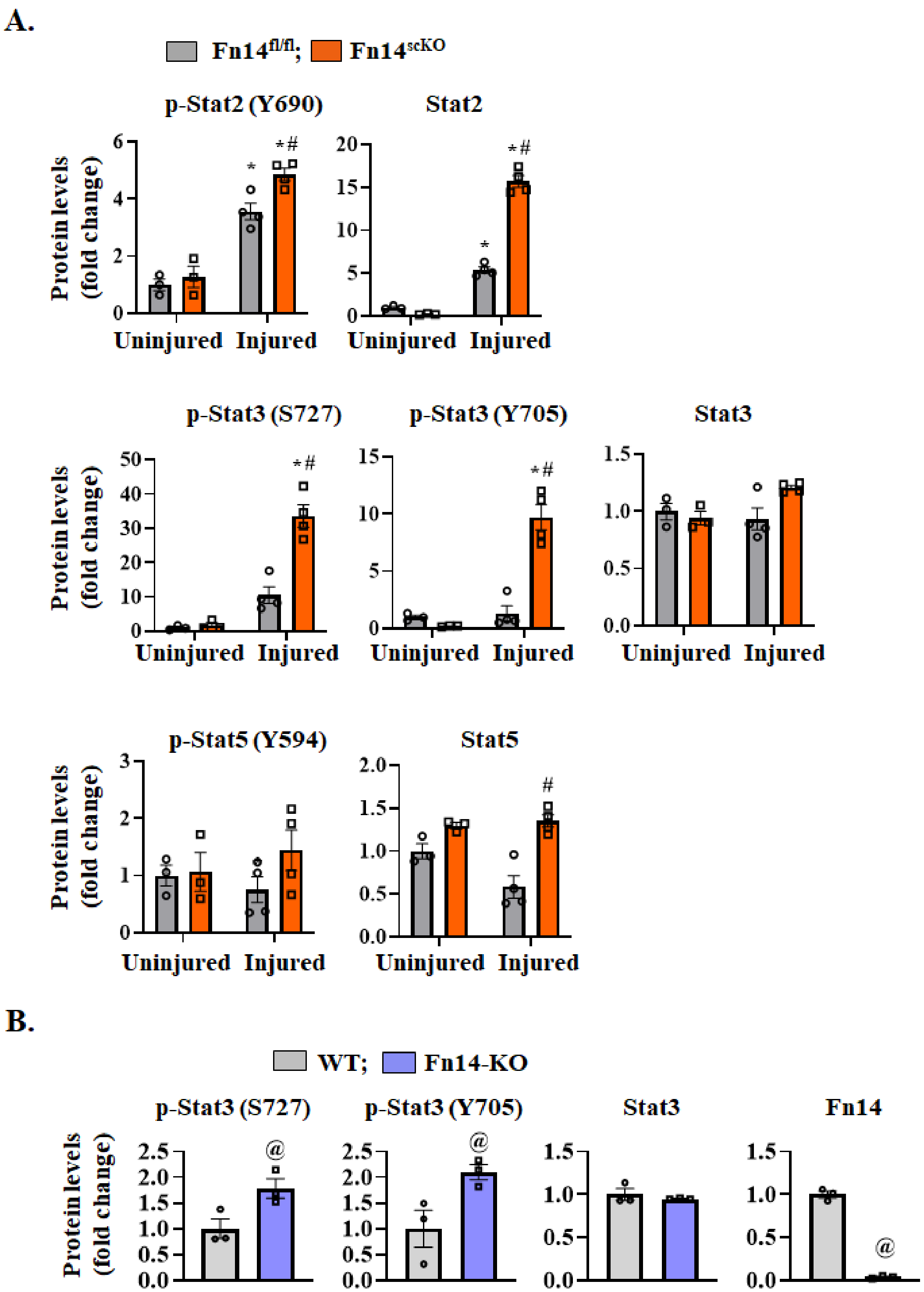


**Supplemental FIGURE 5. Fn14 regulates Stat signaling in injured muscle and cultured myoblasts. (A)** Densitometry analysis of immunoblots (shown in main Figure 7B) demonstrates fold change in the levels of various phosphorylated and total Stat2, Stat3, and Stat5 protein in uninjured and 5d-injured TA muscle of $Fn14^{fl/fl}$ and $Fn14^{scKO}$ mice. n= 3-4 mice in each group.

**(B)** Fold change in the levels of phosphorylated and total Stat3 protein in WT and Fn14-KO cultured myoblasts. n= 3 biological replicates in each group. All data are presented as mean ± SEM. *$p \leq 0.05$, values significantly different from contralateral uninjured muscle of $Fn14^{fl/fl}$ or $Fn14^{scKO}$ mice; #$p \leq 0.05$, values significantly different from corresponding injured TA muscle of $Fn14^{fl/fl}$ mice analyzed by two-way ANOVA followed by Tukey's multiple comparison test. $^{@}p \leq 0.05$, values significantly different from WT cultures analyzed by unpaired Student *t* test.

**Supplemental Table 1**. List of primers used for PCR/qRT-PCR analysis.

| **Name** | **Forward primer (5'-3')** | **Reverse primer (5'-3')** |
|---|---|---|
| Fn14 | AAGTGCATGGACTGCGCTTCTT | GGAAACTAGAAACCAGCGCCAA |
| Pax-7 | CAGTGTGCCATCTACCCATGCTTA | GGTGCTTGGTTCAAATTGAGCC |
| Myod1 | TGGGATATGGAGCTTCTATCGC | GGTGAGTCGAAACACGGATCAT |
| Myh3 | ACATCTCTATGCCACCTTCGCTAC | GGGTCTTGGTTTCGTTGGGTAT |
| Myog | CATCCAGTACATTGAGCGCCTA | GAGCAAATGATCTCCTGGGTTG |
| Notch1 | CAGGAAAGAGGGCATCAG | AGCGTTAGGCAGAGCAAG |
| Notch2 | GCAGGAGCAGGAGGTGATAG | GCGTTTCTTGGACTCTCCAG |
| Notch3 | GTCCAGAGGCCAAGAGACTG | CAGAAGGAGGCCAGCATAAG |
| Jagged1 | AACAAAGCTATCTGCCGACAGG | GGCTGATGAGTCCCACAGTAATTC |
| Jagged2 | TTGGTGGCAAGAACTGCTCAGT | GCTGTCACAGATGCAGGAGAAGTT |
| Dll1 | ACTGTACTCACCATAAGCCGTGCA | TCAGCTCACAGACCTTGCCATAGA |
| Dll4 | CACTTGCCACGATCTGGAGAAT | TGCCCACAAAGCCATAAGGA |
| Hes1 | GCACAGAAAGTCATCAAAGCC | TTGATCTGGGTCATGCAGTTG |
| Hes6 | GCCGGATTTGGTGTCTACAT | TCCTGAGCTGTCTCCACCTT |
| HeyL | CAGATGCAAGCCCGGAAGAA | ACCAGAGGCATGGAGCATCT |
| Hey1 | TGAATCCAGATGACCAGCTACTGT | TACTTTCAGACTCCGATCGCTTAC |
| β-actin | CAGGCATTGCTGACAGGATG | TGCTGATCCACATCTGCTGG |

**Supplemental Table 2**. List of antibodies used for Western blot and Immunofluorescence.

| **Antibody** | **Source and Catalog no.** | **Analysis** |
|---|---|---|
| Monoclonal mouse-anti-Pax7 | DSHB # PAX7 | WB/IF |
| Monoclonal mouse-anti-MyoD | Santa Cruz Biotechnology # 377460 | WB/IF |
| Monoclonal mouse-anti-Myogenin | DSHB # F5D | WB/IF |
| Polyclonal TWEAK Receptor/Fn14 | Cell Signaling Technology # 4403 | WB |
| Monoclonal rabbit-anti-GAPDH | Cell Signaling Technology # 2118 | WB |
| Monoclonal rabbit-anti-Cleaved Notch1 | Cell Signaling Technology # 4147 | WB |
| Polyclonal goat-anti-Notch1 | Santa Cruz Biotechnology # 6014 | WB |
| Polyclonal rabbit-anti-phospho-STAT2(Y690) | Cell Signaling Technology # 4441 | WB |
| Monoclonal rabbit-anti-STAT2 | Cell Signaling Technology # 72604 | WB |
| Polyclonal rabbit-anti-phospho-STAT3 (S727) | Cell Signaling Technology # 9134 | WB |
| Monoclonal rabbit-anti-phospho-STAT3 (Y705) | Cell Signaling Technology # 9145 | WB |
| Monoclonal rabbit-anti-STAT3 | Cell Signaling Technology # 30835 | WB |
| Monoclonal rabbit-anti-phospho-STAT5 (Y694) | Cell Signaling Technology # 4322 | WB |
| Monoclonal rabbit-anti-STAT5 | Cell Signaling Technology # 94205 | WB |
| Monoclonal mouse-anti-CD266 (ITEM-4-Fn14) | e-Bioscience # 14-9018-82 | IF/Flow |
| Rat anti-mouse IgG2b FITC | Invitrogen # 11-4220-82 | Flow |
| PE Rat Anti-mouse-anti- CD31 | BD Pharmingen # 553373 | Flow |
| Monoclonal Rat Anti-mouse-anti-CD45 PE | Invitrogen # 12-0451-82 | Flow |
| Monoclonal Rat Anti-mouse-anti-Ly-6A (Sca-1) PE | e-Bioscience # 12-5981-82 | Flow |
| Monoclonal Rat Anti-mouse-anti-Ter119 PE | e-Bioscience # 12-5921-82 | Flow |
| Monoclonal mouse-anti-α7-integrin APC | Miltenyi Biotec # 130-102-717 | Flow |
| Monoclonal mouse-anti-Myosin heavy chain (embryonic) | DSHB # F1.652 | WB/IF |
| Polyclonal rabbit-anti-Laminin | Sigma # L9393 | IF |
| Polyclonal rabbit-anti-Dystrophin | Abcam # 15277 | IF |
| Goat anti-Mouse IgG1 Alexa Fluor 568 | Life Technologies # A21124 | IF |
| Goat anti-Mouse IgG2 Alexa Fluor 594 | Life Technologies # A211135 | IF |
| Goat anti-Rabbit IgG Alexa Fluor 488 | Life Technologies # 11034 | IF |
| Donkey anti-Rabbit IgG Alexa Fluor 555 | Invitrogen # A31572 | IF |

**Supplemental Table 3.** Mean TPM values for WT myoblasts incubated in growth medium.

| Gene name | Mean TPM values | Gene name | Mean TPM values |
|---|---|---|---|
| Ccng1 | 621.1946774 | Lbh | 249.6727904 |
| Cxadr | 35.79813652 | Lif | 2.59100748 |
| Calcrl | 12.72959725 | Ncoa3 | 9.162479514 |
| Cdk19 | 24.19957131 | Notch1 | 22.85663415 |
| Ccnd1 | 330.2630867 | Notch2 | 37.60329254 |
| Ccnd2 | 104.5287759 | Pax3 | 4.398175455 |
| Hgf | 8.794593864 | Six2 | 27.12187878 |
| Gata6 | 1.084727413 | Sox9 | 35.29842376 |
| Fgfr2 | 0.948072161 | Tead3 | 10.02624689 |
| Apc2 | 1.818422341 | Tgfbr3 | 1.548204592 |
| Enpp1 | 0.989866001 | Il6 | 3.397958934 |
| Nkx2-5 | 0.326197706 | Il15 | 0.419057697 |
| Agtr1a | 1.034336915 | Cxcl10 | 17.12602464 |
| Ccn3 | 4.104003524 | Pdgfb | 4.03449691 |
| Cav2 | 8.514190025 | Camk2a | 5.321223919 |
| Cdkl5 | 0.551536088 | Camk2b | 0.897912176 |
| Ednra | 3.094047633 | Ifnb1 | 0 |
| Cdkn2d | 37.22444052 | Ifna4 | 0 |
| E2f4 | 60.58188387 | Mmp2 | 164.1831704 |
| Cdkn1a | 587.4721457 | Mmp9 | 0.485111336 |
| Igfbp3 | 131.4441299 | Aldh1a1 | 0.545122263 |
| Csrp3 | 28.55942468 | Birc5 | 72.57266062 |
| Cxcl9 | 0.128365652 | Socs2 | 61.92687082 |
| Neu2 | 0.49052196 | Socs3 | 17.30150569 |
| Smyd1 | 66.58618927 | Socs5 | 38.53755219 |
| Lmod3 | 79.02987886 | Sema3a | 36.75054031 |
| Myf6 | 14.00136893 | Sema3e | 95.10299035 |
| Cav3 | 108.0811797 | Fgf7 | 29.93319757 |
| Myog | 1319.010602 | Vegfd | 2.329139809 |
| Rbm24 | 138.0515323 | Il12a | 7.134481129 |
| Hopx | 9.411419736 | Mdm2 | 129.1157167 |
| Fdps | 31.01994176 | Adamts1 | 17.27912129 |
| Olfm2 | 0.642781463 | Cx3cl1 | 16.3392645 |
| Mef2c | 18.38525838 | Cxcl12 | 57.29570539 |
| Ripor2 | 2.677810301 | Dock4 | 1.020768725 |
| Hif1an | 25.38894483 | Sirt1 | 19.41460565 |
| Akap6 | 9.931814853 | Adora2b | 29.72144694 |
| Dpf3 | 1.514276075 | Aqp1 | 23.6756034 |
| Fgfr1 | 78.48068766 | Ackr3 | 93.70005894 |
| Hmga2 | 122.8227459 | Epha1 | 60.8916724 |
| Jmjd1c | 25.22217496 | Epha4 | 1.221308232 |
|  |  | Nedd9 | 12.21753631 |
